# Designed architectural proteins that tune DNA looping in bacteria

**DOI:** 10.1101/2021.06.30.450240

**Authors:** David Tse, Nicole A. Becker, Robert T. Young, Wilma K. Olson, Justin P. Peters, Tanya L. Schwab, Karl J. Clark, L. James Maher

## Abstract

Architectural proteins alter the shape of DNA, often by distorting the double helix and introducing sharp kinks that relieve strain in tightly-bent DNA structures. Here we design and test artificial architectural proteins based on a sequence-specific Transcription Activator-like Effector (TALE) protein, either alone or fused to a eukaryotic high mobility group B (HMGB) DNA-bending domain. We hypothesized that TALE protein binding would stiffen DNA to bending and twisting, acting as an architectural protein that antagonizes the formation of small DNA loops. In contrast, fusion to an HMGB domain was hypothesized to generate a targeted DNA-bending architectural protein that facilitates DNA looping. We provide evidence from *E. coli* Lac repressor gene regulatory loops supporting these hypotheses in living bacteria. Both data fitting to a thermodynamic DNA looping model and sophisticated molecular modeling support the interpretation of these results. We find that TALE protein binding inhibits looping by stiffening DNA to bending and twisting, while the Nhp6A domain enhances looping by bending DNA without introducing twisting flexibility. Our work illustrates artificial approaches to sculpt DNA geometry with functional consequences. Similar approaches may be applicable to tune the stability of small DNA loops in eukaryotes.

## INTRODUCTION

Duplex DNA contains the genetic information of cellular organisms. DNA is also one of the stiffest biopolymers (1). Overcoming DNA stiffness is important for various biological processes such as compaction of viral DNA into bacteriophage heads (2) and formation of small DNA loops (3). The importance of DNA looping cannot be understated. Distant sites separated across the genome are brought together in transcriptional gene regulation and recombination (4,5). In some cases, multiple genes are transcribed together through a shared RNA polymerase in a transcription “factory” (6). Control of these processes might be achieved through the artificial manipulation of DNA stiffness. Here we explore the design and testing of sequence-specific architectural DNA binding proteins to tune DNA looping over short distances in bacteria where resistance of DNA to bending and twisting is dominant. We achieve this by engineering Transcription Activator-like Effector (TALE) proteins with or without fusion to a eukaryotic high mobility group B (HMGB) domain capable of inducing sharp DNA bending. Part of this work emulates with designed proteins features of tight DNA loop enhancement by natural sequence-nonspecific architectural proteins (7,8).

TALEs were discovered as virulence factors in phytopathogenic bacteria from the genus *Xanthomonas*. These bacteria utilize a type III secretion system to inject TALEs into plant cells to activate genes that apparently facilitate infection (9). TALEs bind DNA in a remarkable sequence-specific manner through well-characterized repeats that consist of 33-35 amino acid modules differing only at positions 12 and 13 (the repeat variable diresidue, RVD). Each diresidue determines the base pair recognition specificity of each corresponding TALE module (10). Crystal structures of TALE binding to DNA show the direct contact between the side chain of residue 13 and the target DNA base, thereby contributing to DNA-binding specificity (11). TALEs found in nature range from 2 to 34 repeats, with the majority being 16-20 repeats in length (9). The optimum length of engineered TALE repeats is 16-25, corresponding to the median length of naturally-occurring TALEs (12).

While no prior studies have addressed the impact of TALE binding on DNA flexibility, inspection of X-ray crystal structures suggests that the TALE amino acids fully engage the DNA major groove (11,13). We hypothesized that this engagement would limit DNA bending and twisting motions, effectively stiffening the DNA polymer. Such stiffening could have the effect of increasing the energy of DNA looping by constraining DNA bending and twisting to a smaller number of base pairs within the loop. In this way, TALEs might serve as artificial architectural DNA binding proteins that tune DNA looping by making it more expensive.

Eukaryotic high mobility group (HMG) non-histone proteins were first isolated almost fifty years ago and named for their electrophoretic mobility in denaturing polyacrylamide gels (14). Many such proteins bind to DNA in a sequence-nonspecific manner. HMG proteins are subdivided into three superfamilies (15), HMGB, HMGN, and HMGA. These proteins act as accessory architectural factors involved in nucleosome and chromatin modulation (16). Architectural proteins are also believed to regulate other nuclear activities including transcription, replication, and DNA repair (17–21). Some HMGB proteins contain two tandem HMG box domains preceded or followed by protein segments rich in acidic residues. Each HMG box is composed of three α-helices that fold into an L-shaped conformation and can bind to the minor groove of DNA, accompanied by intercalation of certain side chains with low sequence specificity (22). Introduction of the hydrophobic surface of the HMG box domain into the minor groove of DNA causes an approximately 90° kink (23). Transient sharp bending of DNA amounts to an enhancement in DNA chain flexibility (24). Nhp6A is a single-box HMGB protein from *Saccharomyces cerevisiae* (25). This sequence-nonspecific architectural protein bears structural similarities to each of the two HMG boxes of mammalian HMGB1/2 proteins. In the present study, Nhp6A is fused to a sequence-specific DNA-binding protein to create a novel architectural DNA binding protein that targets a DNA kink to a desired site within a DNA repression loop.

TALEs have been shown to be effective for tethering other functions such as endonucleases (26–28). We therefore hypothesized that TALE-HMGB protein fusions might serve as artificial sequence-specific architectural proteins that deliver sharp DNA bends, facilitating DNA looping. Such designed artificial architectural DNA binding proteins would offer new tools for programing the three-dimensional structure of DNA. We set about to test these ideas in this work.

Our approach is summarized in Fig. 1. We hypothesized that a strongly-expressed reporter gene (Fig. 1A) will be repressed by a small and strained DNA loop (Fig. 1B), and that this repression can be tuned by artificial TALE-based architectural DNA binding proteins that either stiffen the looped DNA leading to derepression (Fig. 1C), or that introduce a kink within the loop, facilitating repression (Fig. 1D). Lac repressor and TALE derivatives (Fig. 1E), are expressed separately from a *lacZ*-based reporter plasmid (Fig. 1F). The impacts of designed artificial architectural DNA binding proteins were assessed in sets of experiments where the architectural protein was placed at a fixed position relative to the downstream operator as loop sized was systematically changed (Fig. 1G), or systematically moved within a DNA loop of fixed length (Fig. 1H).

**Fig. 1.**
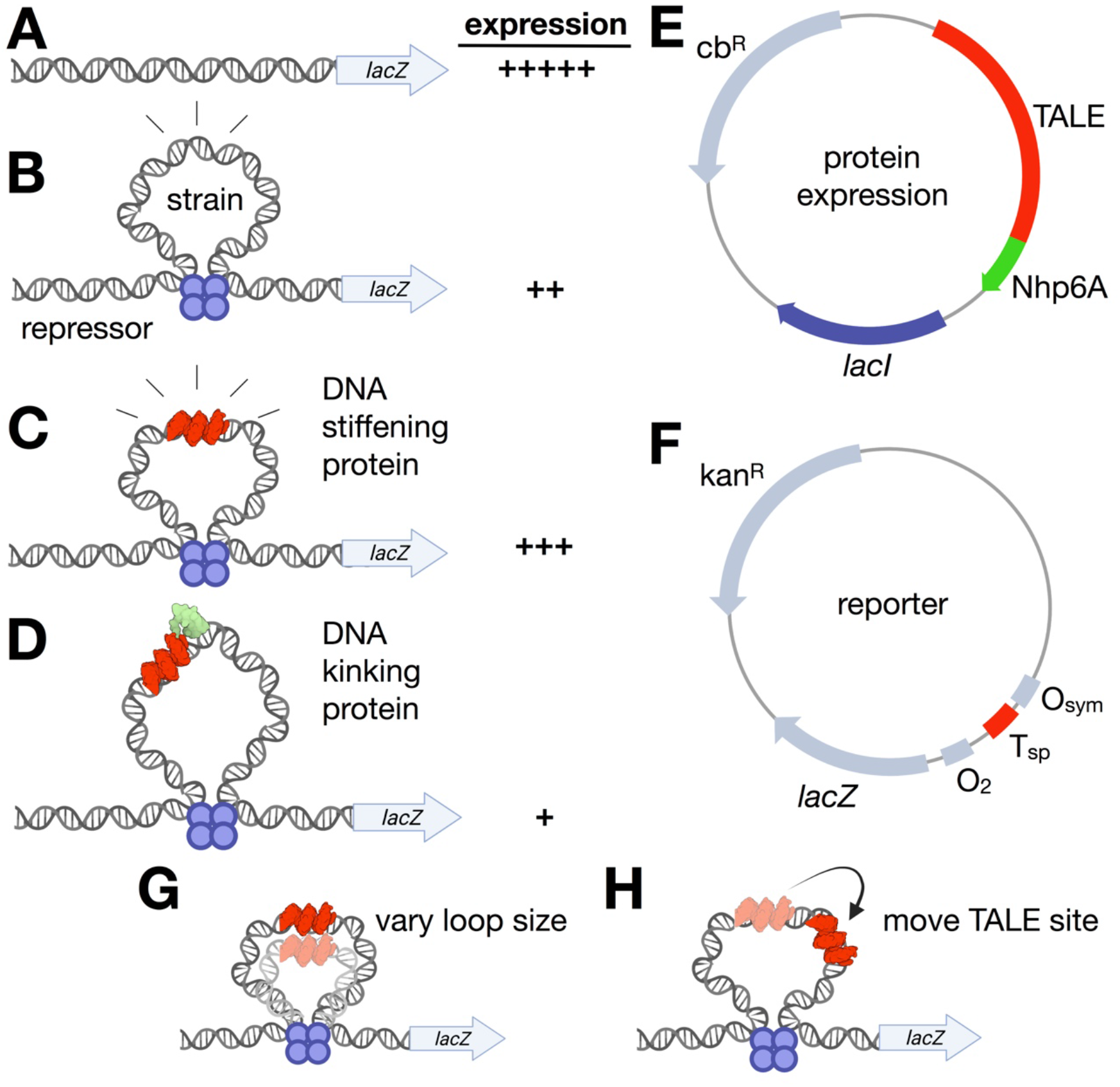
Concepts addressed in this work. A. Reporter construct in the absence of lacI-mediated looping is predicted to be strongly expressed. B. Small, strained DNA loops anchored by lacI should substantially repress the promoter of the reporter construct. C. It is hypothesized that a TALE protein targeted to the lacI loop will serve as an artificial architectural DNA binding protein that stiffens the bound DNA segment to bending and twisting, limiting deformations to unbound segments of the loop, and thereby shifting the equilibrium away from looping and causing derepression. D. It is hypothesized that endowing the site-specific TALE protein with a sequence-nonspecific Nhp6A DNA-kinking domain will relieve bending strain, improving repression by lacI-mediated looping. E. Schematic illustration of protein expression plasmid providing TALE protein with or without Nhp6A fusion, and/or lacI. F. Schematic illustration of experimental reporter construct to evaluate detailed effects of artificial sequence-specific architectural proteins. G. Scheme for reporter constructs that vary loop size with a fixed TALE position. H. Scheme for reporter constructs that vary TALE position for a fixed loop size.

## MATERIALS AND METHODS

### DNA looping reporter constructs

DNA looping reporter constructs were based on plasmid pJ2656 containing the *lac* uv5 promoter and a single proximal O_2_ operator. Molecular cloning was used to install a 15-base-pair specific TALE recognition sequence (T_sp_) (Supplemental Table S1) – top strand: tTCATGTTATAACGGA to generate plasmid pJ2721 containing the T_sp_ site upstream of the proximal O_2_ operator as the basis for Series 1 constructs. The 5′ invariant thymine residue of the TALE target is shown in lower case. Various oligonucleotide duplexes containing the O_sym_ operator were then cloned to generate pJ2721-pJ2737 constructs (Supplemental Table S2, Supplemental Fig. S1) with spacings of 131.5 bp - 146.5 bp (measured center-to-center between O_2_ and O_sym_ operators). Plasmid pJ2748 was created from pJ2721 using molecular cloning upstream of the T_sp_ recognition sequence to serve as the basis for Series 2 constructs. Duplexes adjust the position of the T_sp_ recognition sequence in base pair increments in the context of a constant 142.5 bp DNA loop length (Supplemental Table S3, Supplemental Fig. S2).

### Expression of DNA binding proteins

Plasmid pJ2652 encodes the specific TALE (sp TALE) protein recognizing the T_sp_ sequence. Plasmid pJ2654 encodes a different TALE (ns TALE) protein recognizing a different sequence (T_ns_ – top strand: tTACAAGTGGCTCATT) (Supplemental Table S1) (29). DNA encoding the yeast Nhp6A protein was amplified from plasmid J2472 and this coding sequence was used create TALE-Nhp6A fusions (Supplemental Table S4). The *lacI* gene encoding Lac repressor was amplified from plasmid pJ2179 and installed downstream from the TALE coding sequence in some cases.

Protein expression plasmids were transformed into electrocompetent bacterial strain FW102 (30) and transformants selected on LB agar plates containing streptomycin and carbenicillin. The resulting bacteria were made electrocompetent and transformed with appropriate reporter plasmids with selection on LB agar plates containing streptomycin, carbenicillin, and kanamycin.

### Western blotting

Bacterial strains were grown in 10 mL LB medium containing appropriate antibiotics overnight at 37 °C with aeration. Saturated overnight culture (200 μL) was used to seed a 20-mL subculture on the next day and grown for 2-3 h at 37 °C with aeration. When the OD_600_ of the subculture reached 0.4-0.6, 6 mL of the culture was pelleted by centrifugation and resuspended in 200 μL 1× NuPAGE MES SDS running buffer (Thermo). Cells were lysed by sonication and clarified by centrifugation. Protein was quantified using a BCA protein assay kit (Thermo). Protein (6 μg) was subjected to electrophoresis through a NuPAGE 10% Bis-Tris precast gels (Thermo) at 125 V for 2 h. Gels were then blotted onto PVDF membranes (Bio-Rad) using an XCell II Blot Module (Thermo) in 1× NuPAGE transfer buffer (Thermo) containing 20% methanol. Blotted membrane was briefly washed in TBST (50 mM Tris-HCl, pH 7.4, 150 mM NaCl, 0.1% Tween 20), treated with blocking buffer (5% dry milk and 1% BSA in TBST) for 1 h at room temperature, incubated with primary antibody overnight at 4 °C in TBST with gentle rocking. Membranes were washed with TBST three times for 10 min at room temperature and then incubated with secondary antibody in blocking buffer for 1 h at room temperature with gentle rocking. Membranes were analyzed with an Amersham Typhoon laser-scanner platform (Cytiva) using IRshort and IRlong methods. Uncropped images are shown in Supplemental Fig. S3.

#### Primary antibodies for western blotting were as follows

mouse anti-AcV5 (1:2000, ab49581, abcam); rabbit anti-*E. coli* Lac repressor (1:2000, C60143, LSBio). Secondary antibodies for western blotting were as follows: IRDye 800CW goat anti-mouse IgG (1:10000, 926-32210, Li-cor); IRDye 680LT goat anti-rabbit IgG (1:10000, 926-68021, Li-cor).

### *E. coli* β-*galactosidase reporter assay*

Quantitation of DNA looping was accomplished by measurement of *β*-galactosidase activity in bacterial extracts as previously described (31). To establish ideal levels of Lac repressor activity, bacterial extracts were assayed after cell growth with or without 100 μM IPTG. *β*-galactosidase reporter activity is presented in Miller units (MU). Repression level (*RL*) is calculated according to equation 1:

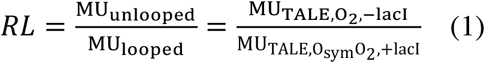

where TALE indicates the presence of any appropriate TALE protein. The normalized repression level (*RL_n_*) is given by equation 2:

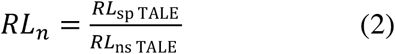

where sp TALE indicates the presence of the specific TALE protein with the target T_sp_ sequence in the reporter constructs, and ns TALE indicates the presence of the ns TALE protein with no binding site present in the reporter system. Data are provided in Supplemental Tables S5 and S6.

### Data fitting to thermodynamic DNA looping model

Curve fitting to a thermodynamic model of *lac* promoter repression was performed using non-linear least-squares refinement to each set of *RL* data with four adjustable parameters for the variable O_sym_–O_2_ series (Series 1) and three adjustable parameters for the variable TALE–O_2_ series (Series 2), as described below (and see Table 1). The thermodynamic model that relates gene expression and spacing of operator sequences has been previously described in (31–33). Fit parameters give insight into the physical properties of the nucleoprotein loops.

**Table 1.** Thermodynamic model fit parameters

| | $O_{\text{sym}} - O_2$ spacings | | | |
| --- | --- | --- | --- | --- |
| Parameter | sp TALE | sp TALE-Nhp6A | ns TALE | ns TALE-Nhp6A |
| $hr$ (bp/turn) | $10.96 \pm 0.03$ | $10.98 \pm 0.07$ | $11.06 \pm 0.37$ | $11.13 \pm 0.51$ |
| $C_{\text{app}}$ ($\times 10^{-19}$ erg-cm) | $0.66 \pm 0.01$ | $0.99 \pm 0.01$ | $0.44 \pm 0.01$ | $0.31 \pm 0.01$ |
| $K_{\text{max}}$ | $11.38 \pm 0.06$ | $63.99 \pm 0.86$ | $33.56 \pm 0.50$ | $29.84 \pm 0.26$ |
| $sp_{\text{optimal}}$ (bp) | $142.57 \pm 0.02$ | $142.80 \pm 0.04$ | $144.03 \pm 0.17$ | $143.26 \pm 0.41$ |
| | TALE – $O_2$ spacings | | | |
| $^{\text{TALE}}C_{\text{app}}$ ($\times 10^{-19}$ erg-cm) | $0.34 \pm 0.01$ | $0.36 \pm 0.01$ | $0.25 \pm 0.01$ | $0.24 \pm 0.01$ |
| $^{\text{TALE}}K_{\text{Smax}}$ | $15.62 \pm 0.12$ | $49.66 \pm 0.97$ | $42.03 \pm 1.03$ | $38.61 \pm 0.99$ |
| $^{\text{TALE}}sp_{\text{optimal}}$ (bp) | $97.89 \pm 0.04$ | $94.79 \pm 0.08$ | $92.95 \pm 0.15$ | $93.18 \pm 0.17$ |

All parameters are +LacI in the presence of 100 $\mu\text{M}$ IPTG and are presented with a 95% confidence interval.

The thermodynamic model is based on the premise that promoter repression is sensitive only to the occupancy of the downstream *lac* O_2_ operator at equilibrium (34–36). The extent of promoter repression is modeled by evaluating the distribution of possible states of this operator. If a singly-bound repressor exists at O_2_ (“single bound”), repression is modest because transcription elongation can, with high probability, disrupt the bound repressor. In contrast, repressor bound to the downstream operator by virtue of DNA looping from the strong upstream O_sym_ operator (“specific looping”) has the potential to cause more complete promoter repression through the mechanisms in question here.

In the variable O_sym_–O_2_ reporter constructs (Series 1), the fraction of proximal operator bound by repressor as a function of DNA operator-operator length is modeled with four adjustable parameters evaluating the distribution of possible states of the proximal operator through a partition function for the system (31,32,37,38). In this context, *hr* is the DNA helical repeat, *C*app is the apparent torsional modulus of the DNA loop, *sp*_optimal_ is the optimal spacing between operators (in base pairs) where *sp* is the actual spacing for a given construct, and *K*_max_ is the equilibrium constant for the formation of a specific loop with optimal phasing.

In the variable TALE–O_2_ reporter constructs (Series 2), the distance between the downstream and upstream operators is held constant at 142.5 bp. The face of the helix occupied by various sp TALE proteins is then systematically changed in a manner that potentially influences loop stability: if TALE binding increases DNA strain, the loop is destabilized by decreased anchoring from repressor or decreased engagement of RNA polymerase. If TALE binding decreases DNA strain, the loop is stabilized (“stabilized loop”) and the repressed state is favored. Here, there are three adjustable parameters, since *hr* remains unchanged from fitting in the previous series. ^TALE^*C*_app_ is the apparent torsional modulus of the DNA loop, ^TALE^*sp*_optimal_ is the optimal spacing between the TALE binding site and O_2_ (in base pairs), and ^TALE^*K*_Smax_ is the equilibrium constant for the formation of a stabilized loop. The thermodynamic model that describes gene expression in the context where the loop size was held constant has been described (39).

### Molecular modeling

The configurations of DNA chains capable of looping between the headpieces of the Lac repressor assembly were obtained using a procedure that optimizes the energy of a collection of base pairs, in which the first and last pairs are held fixed (40). The DNA is described at the level of base-pair steps using six rigid-body parameters to specify the arrangements of successive base pairs — three angles (tilt, roll, twist) describing the orientation of successive base-pair planes and three translational components (shift, slide, rise) along the vector joining successive base-pair centers (41–43). The base pairs in contact with the repressor and TALE-Nhp6A constructs are held fixed, with rigid-body parameters assigned values extracted from high-resolution structures (11,25,44–47) (see Supporting Information). The protein-free steps are subject to a potential that allows for elastic deformations of DNA from its equilibrium structure (48). The steps are assigned the elastic properties of an ideal, inextensible, naturally straight double helix, with bending deformations consistent with the persistence length of mixed-sequence DNA (49), fluctuations in twist compatible with the topological properties of DNA minicircles (50,51), and a variable helical repeat. These features are expressed at the base-pair level in terms of an equilibrium rest state with null values of all rigid-body parameters other than twist and rise and a set of elastic constants impeding deviations of parameters. The twist is assigned a reference value of 360°/*n*, where *n* is the assumed number of base pairs per turn, and the rise a value of 3.4 Å, the distance between the planes of successive base pairs. The energy is increased by ½ *k*_B_*T* by a change in twist of 4.1°, a bending deformation of 4.8°, or a translational move of 0.02 Å, where *k*_B_ is the Boltzmann constant and *T* the absolute temperature. Stabilizing interactions between protein and DNA are not considered. Configurations with steric overlaps are discarded.

In the absence of knowledge of the directions in which the DNA operators attach to the arms of the repressor, each operator is placed in two orientations on the protein-binding headpieces, yielding four distinct loops — two termed A_1_ and A_2_ with the 5′-3′ directions of the bound fragments running in nearly opposing (antiparallel) directions and two termed P_1_ and P_2_ with the fragments running in the same (parallel) direction (52). Moreover, each of these loops includes configurations from two competing structural families, with similar, albeit out-of-phase, dependencies on chain length (53,54), leading to eight potential spatial forms 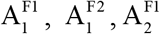, etc. distinguished by the family (F1, F2) and connectivity (1, 2) of the loop. The leading strand of the O_sym_ operator at the start of the loop progresses toward the central axis of the protein assembly in configurations with a connectivity of 1 and toward the outside in those with a connectivity of 2.

Optimized structures of DNA loops of increasing chain length are obtained from the configurations of previously determined protein-free loops with 92-bp center-to-center operator spacing (54). Base pairs are added one at a time by assigning the coordinates of an arbitrary base pair in an existing structure to a new residue and then minimizing the energy of the enlarged system under the same end-to-end constraints. Proteins are introduced with a ramping procedure that gradually changes the rigid-body parameters of base-pair steps at the desired binding site from those of deformable, protein-free DNA. Each ramping step is accompanied by an optimization, which freezes the partially deformed protein-bound segment while reconfiguring the DNA loop.

Looping propensities, or *J* factors, are estimated from the sums of the statistical weights, *i.e*., the Boltzmann factors, of the eight energy-optimized configurations of a loop of given chain length in the presence or absence of a specifically bound TALE-Nhp6A construct. If a DNA loop is short enough, the lowest energy states mirror the looping propensities and modes of chain attachment captured in direct simulations of linear chain molecules subject to the same spatial constraints (53,55). The preferred modes of looping are measured in terms of the fractional contributions *f_i_* of each specific loop type *i* to the *J* factor, i.e.,

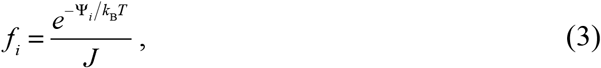

where Ψ_*i*_ is the total elastic energy of the optimized loop and *J* is evaluated over the eight looped states.

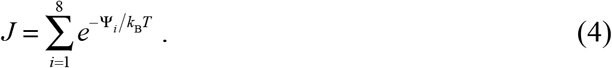

## RESULTS AND DISCUSSION

### TALE-based architectural protein design

TALE and TALE-Nhp6A (Fig. 2 and Supplemental Table S4) were created by replacing the FKBP (F36M) in TALE dimer constructs (29) with a stop codon or Nhp6A domain (Fig. 2C). The *lac* operon looping system is anchored through the binding of Lac repressor tetramer at O_sym_ and O_2_ operators (Fig. 1B and Supplemental Fig. S1). The *lacI* gene encoding Lac repressor was cloned downstream of the desired TALE coding sequence. In this way, DNA looping was enabled or disabled by including or excluding Lac repressor from the bacterial system.

**Fig. 2.**
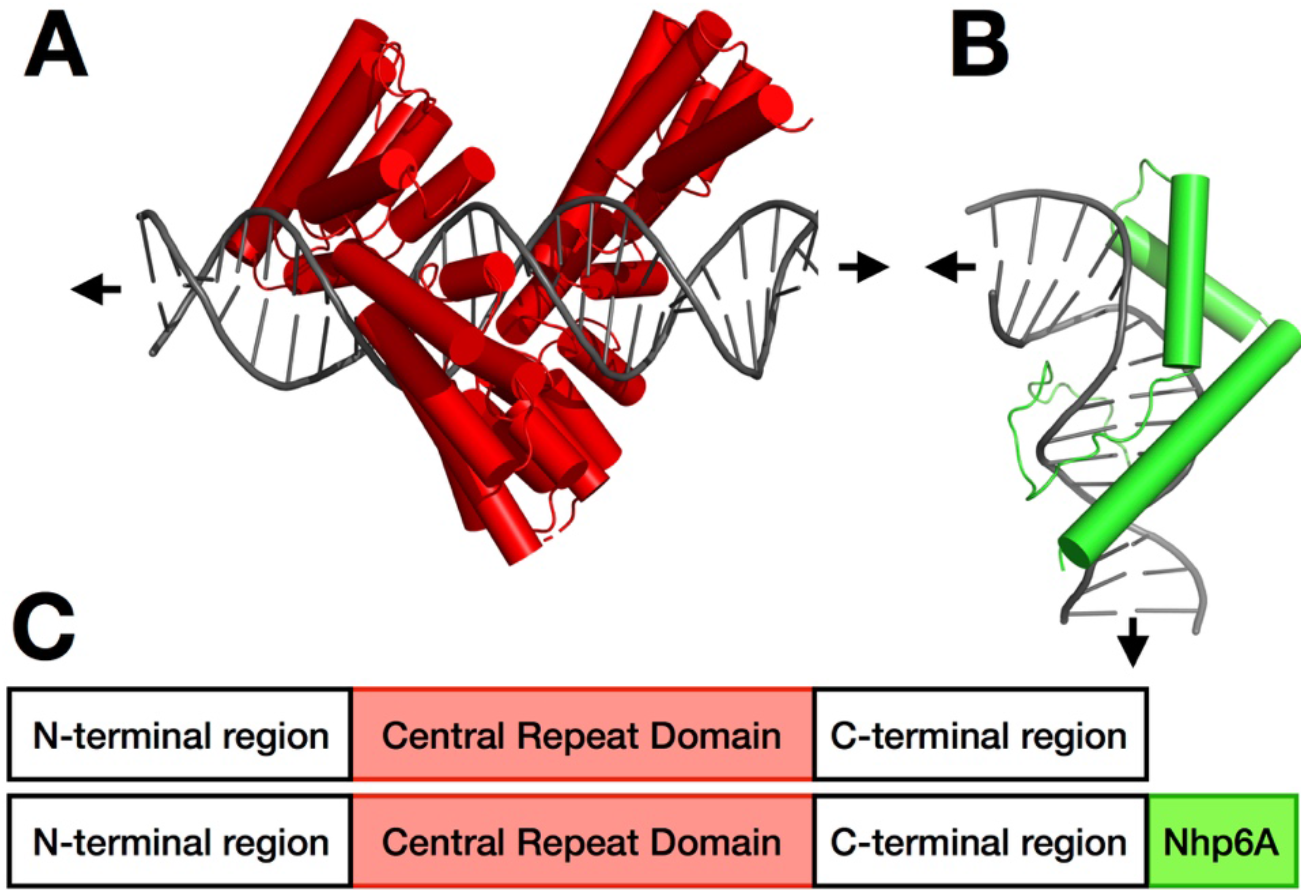
Engineered fusion proteins. A. General TALE structure (red) bound to DNA (grey) based on PBD: 3UGM, but with 15-bp target site as in this work. B. Nhp6A structure (green) bound to DNA (grey) based on PDB: 1CG7. Arrows indicate DNA trajectory. C. Domain structure of fusion proteins.

### Characterization of components

TALE and TALE-Nhp6A (Fig. 2) were expressed in bacterial strain FW102 (30). Western blotting (Fig. 3A) reveals that TALE, TALE-Nhp6A, and Lac repressor proteins are properly expressed at their expected molecular weights. Importantly, expressed levels of TALE proteins and Lac repressor are independent. The DNA binding specificities and affinities of TALE proteins and fusions were assessed using *β-*galactosidase reporter assay (Fig. 3B) where reporter constructs contained T_sp_ or T_ns_ DNA recognition sequences (Supplemental Table S1) in a position corresponding to the proximal operator but in the absence of Lac repressor. Binding of the TALE protein in this position inhibits transcription and subsequent *lacZ* expression. *β*-galactosidase reporter assays sensitively reflect TALE binding affinity, revealing high *lacZ* expression when sp TALE and sp TALE-Nhp6A are expressed with reporter constructs containing the non-cognate proximal T_ns_ TALE recognition sequence (Fig. 3B, black bars at right). Conversely, *lacZ* expression is repressed when sp TALE and sp TALE-Nhp6A are co-expressed with a reporter plasmid containing a cognate proximal T_sp_ recognition sequence (Fig. 3B, black bars at left). Likewise, co-expression of ns TALE and ns TALE-Nhp6A with a reporter construct containing a proximal T_ns_ DNA recognition sequence results in lower *lacZ* expression, though repression is not as strong as for sp TALE and its cognate T_sp_ site (Fig. 3B, grey bars). Interestingly, binding of sp TALE-Nhp6A fusions causes greater repression than the corresponding TALE proteins (Fig. 3B). This could imply a repressive effect of an induced DNA kink at the position of the Nhp6A module, or increase steric occlusion of the promoter by this C-terminal module. Together, these data show that TALE constructs have high binding specificity for their target recognition sequences.

**Fig. 3.**
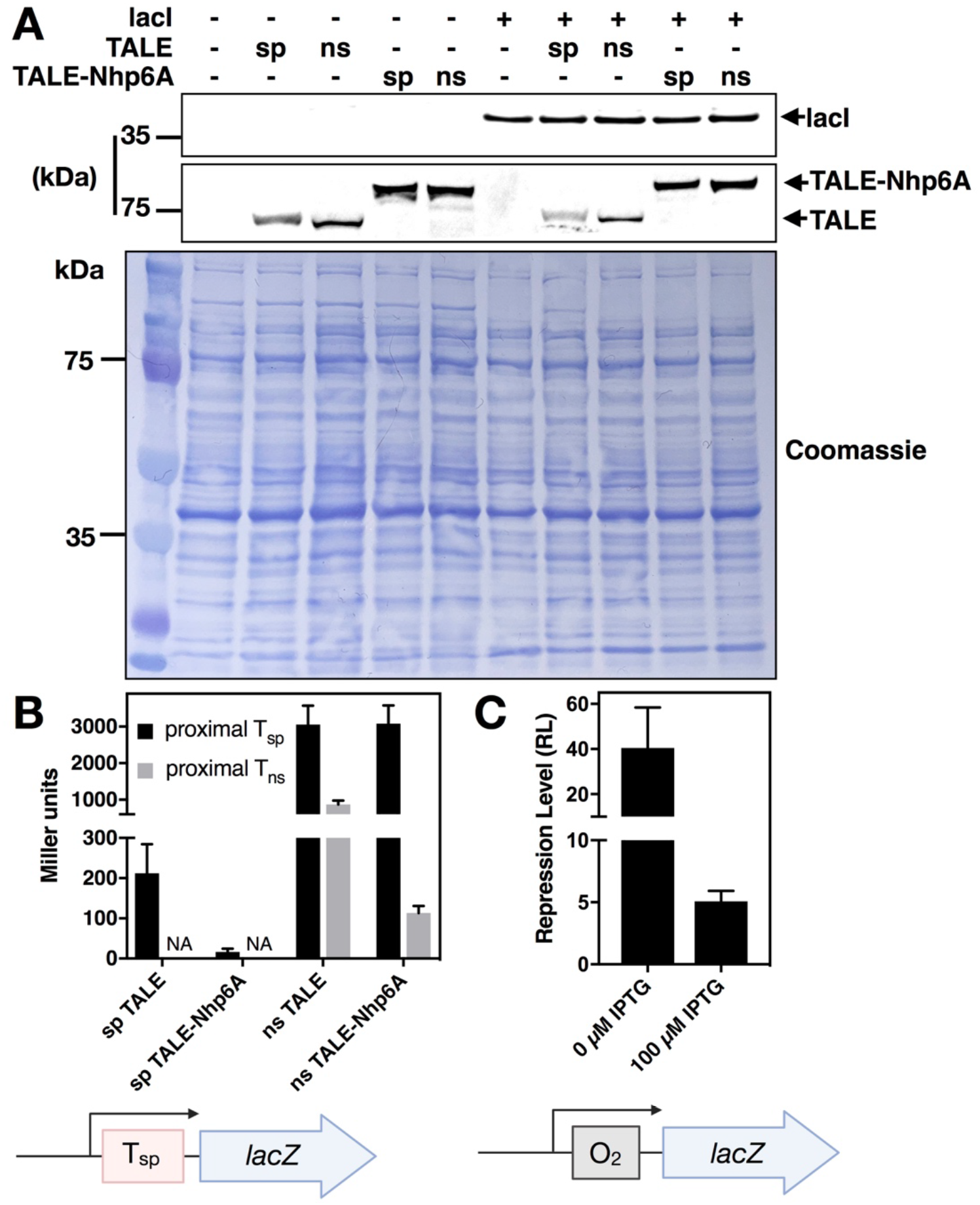
lacI and TALE protein expression, TALE binding affinity and specificity, and lacI tuning with IPTG. A. Western blot analysis of lysates from *E. coli* expressing the indicated genes encoding specific (sp) or nonspecific (ns) TALE (75.4 kDa), or TALE-Nhp6A (86.2 kDa) fusion proteins with or without lacI (38.6 kDa). TALE and TALE-Nhp6A fusion proteins contain an N-terminal AcV5 epitope. B. lacZ expression when the indicated TALE (x-axis) is expressed with the indicated TALE binding sequence at the proximal operator position. C. Desired tuning of effective lacI affinity for test reporter construct with O_2_ in the proximal position.

For convenience, the present assay system employs multi-copy plasmids rather than single-copy episomal constructs as we often study. This results in a higher Lac repressor expression and corresponding greater basal repression from reference reporters carrying a single weak pseudo-palindromic O_2_ operator in the proximal position [~40-fold repression rather than ~4-fold observed for Lac repressor expressed from a single-copy episome (32)]. To compensate and facilitate comparison with this earlier work, Lac repressor affinity was moderated by performing all experiments in the presence of a low concentration (100 μM) of IPTG inducer, bringing *RL* to ~ 5 (Fig. 3C).

### *lac* looping model system

We and others have engineered elements derived from the *lac* operon to create models allowing the study of DNA flexibility in living bacteria and effects of architectural DNA binding proteins on DNA looping (8,37,38). Intrigued by the idea of using designed sequence-specific TALE proteins as artificial architectural proteins to tune DNA looping, we began by designing reporter constructs (Fig. 1, F-G) containing the *lac* O_2_ operator in the proximal position to control *lacZ* transcription by RNA polymerase. Consistent with our past designs, an O_sym_ operator is then installed 131.5 - 146.5 bp (measured center-to-center) upstream from O_2_. The T_sp_ TALE recognition site is placed at a fixed position 85.5 bp upstream to the O_2_ operator (Series 1 constructs; Fig 1G, Supplementary Fig. S1). Upon binding of the Lac repressor tetramer to both operators, a DNA loop is formed. Varying the distance between O_sym_ and O_2_ results in loops that differ in both length and twist strain. In a second set of experiments, the distance between O_sym_ and O_2_ operators was fixed at 142.5 bp (untwisted loop) and the T_sp_ TALE recognition site position was varied from 85.5 to 100.5 bp (measured center-to-center) upstream from the O_2_ operator (Series 2 constructs, Fig. 1H, Fig. 4B, top diagram, Supplementary Fig. S2). This series rotates, at base-pair resolution, the placement of the artificial architectural DNA binding protein on different DNA faces within the DNA loop. In both Series 1 and Series 2 experiments we hypothesized that the effects of artificial architectural proteins would depend on loop geometry and twist strain.

**Fig. 4.**
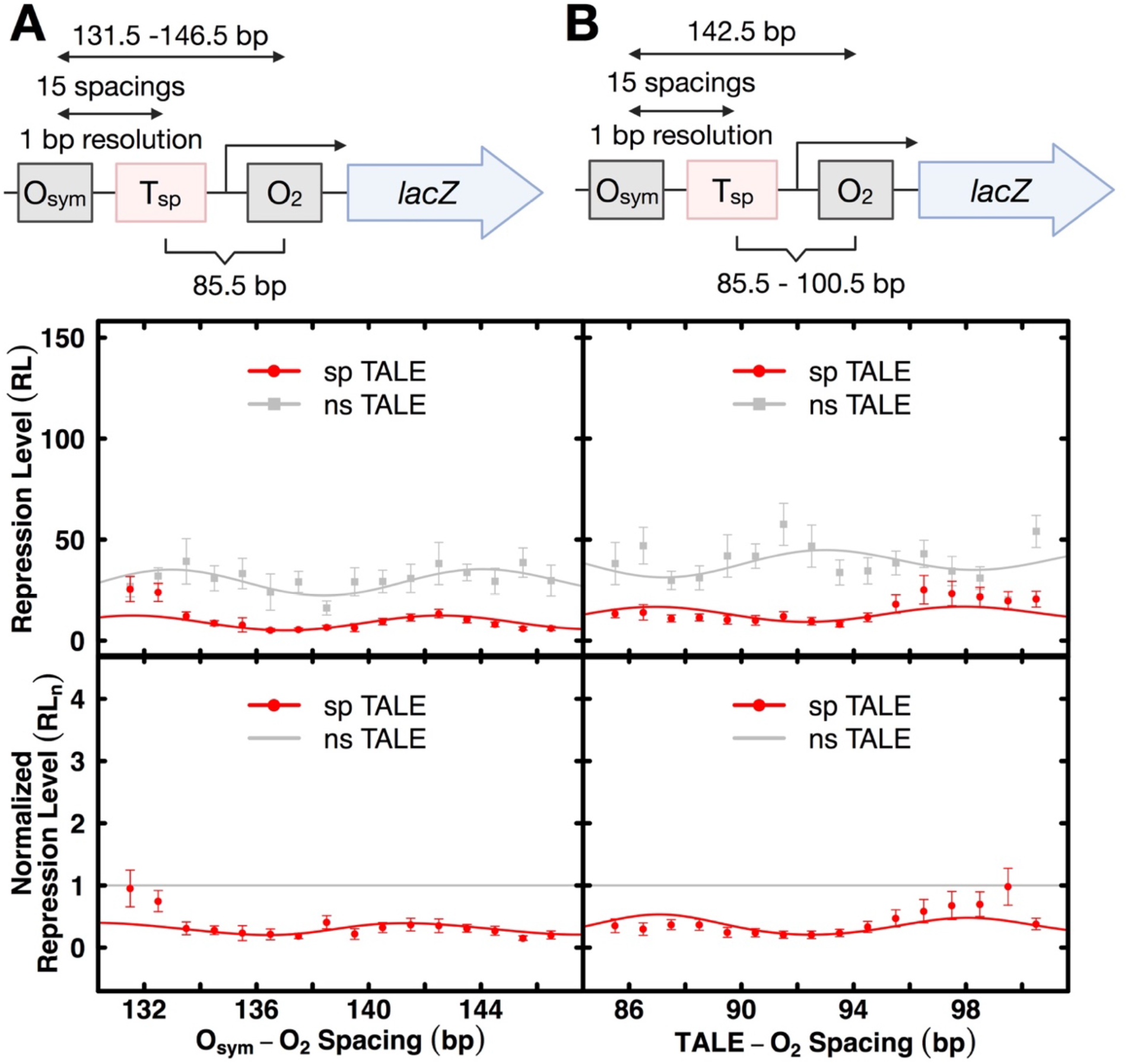
Effects of designed TALE proteins on lacI-mediated DNA looping in living *E. coli*. A. Series 1 construct. The top diagram indicates control element configuration. The upper graph depicts repression level (*RL*) as a function of center-to-center lac operator spacing in the presence of the indicated nonspecific (grey) or specific (red) TALE protein where the TALE site is at a fixed position relative to O_2_ in the presence of lacI and 100 μM IPTG. The lower graph compares nonspecific or specific TALE effects when repression level values are normalized to the corresponding values for nonspecific designed architectural protein (grey line indicates a value of 1.0). B. As in panel (A) except the data were obtained with series 2 constructs illustrated in top diagram. All depicted curve fits are to the thermodynamic model described in methods, with fit parameters reported in Table 1.

### The TALE as a DNA stiffening architectural protein to antagonize looping

TALEs were co-expressed with Series 1 and Series 2 reporter spacing constructs (Fig. 4AB, top diagrams). Series 1 reporter constructs alter operator spacing in 1-bp increments to create repression loops of different lengths and twist strains. Repression is monitored by the repression level parameter (*RL*, Materials and Methods equation 1). It is important to emphasize that *RL* is defined by repression measurements obtained in the presence of sp TALE binding, so any effects of TALE binding on basal promoter function are taken into account by this parameter. In Series 1 reporter constructs (Fig. 4A, upper graph) studied in the presence of ns TALE that cannot bind the loop, repression shows a modest dependence on operator spacing (grey data and fit) as expected for loops of this length, reflecting the expense of DNA twisting. Greatest repression occurs for untwisted loops with operator spacings near 133 bp and 144 bp, indicating an in vivo helical repeat of 11 bp/turn as commonly observed for negatively-supercoiled domains in bacteria. In striking contrast, sp TALE binding within the loop causes global derepression, consistent with the intended antagonism of looping by DNA stiffening (Fig. 4A, upper graph, red data and fit). This effect is emphasized by normalizing to the data for the ns TALE (Fig. 4A, lower graph).

Quantitative thermodynamic modeling data supporting these interpretations are presented in Table 1. The observed helical repeat (*hr*) is ~11 bp/turn, setting the phasing for one periodic *sp*_optimal_ at ~143 bp (~13 helical turn operator separation). The fit values of the specific looping equilibrium constant, *K*_max_, in the presence of ns TALEs are 29.8-33.6, (combined 31.7). Importantly, this value decreases by a factor of ~3 (11.4) for the sp TALE. This result parallels the observed decrease in *RL* values and shows that the sp TALE antagonizes DNA looping. Interestingly, values of the DNA twist constant, *C*_app_, in the presence of ns TALE are 0.31-0.44 (combined 0.37) but in the presence of the sp TALE range from 0.66-0.99, more than twice as large. This striking result indicates that TALE binding creates a large obstacle to both DNA bending and twisting.

Similar results are obtained for Series 2 reporter constructs that incrementally reposition the T_sp_ TALE binding site within an untwisted repression loop where the Lac operators are spaced by 142.5 bp (Fig. 4B, top diagram). Comparing repression data obtained in the presence of a nonspecific vs. specific TALE, derepression is again observed for all positions with sp TALE binding, consistent with an intended DNA stiffening effect that should not depend on helical phasing (Fig. 4B, upper graph). Again, normalization to repression data in the presence of the ns TALE emphasizes this result (Fig. 4B, lower graph). Thus, global derepression is caused by sp TALE binding within the repression loop, consistent with a stiffening effect of sp TALE on DNA bending and twisting (Supplemental Tables S5 and S6). Quantitative data fitting to the thermodynamic model is again shown in Table 1, supporting these interpretations. In the presence of the ns TALE, the DNA twist constant, ^TALE^*C*_app_, is ~0.25, ^TALE^*K*_Smax_ is ~40 and ^TALE^sp_optimal_ is ~93. When compared to sp TALE, ^TALE^*C*_app_ increases to ~0.34 and ^TALE^*K*_Smax_ decreases ~2.6 fold to ~16, again providing evidence that the designed TALE architectural protein stiffens DNA to both bending and twisting, antagonizing looping as hypothesized.

### The TALE-Nhp6A fusion as a DNA bending architectural protein to facilitate looping

We hypothesized that fusion with the sequence-nonspecific yeast Nhp6A HMGB DNA bending protein would allow targeted DNA bending within the repression loop to alter its energetics. Results are presented in Fig. 5, showing an intriguing phasing dependence of TALE-Nhp6A on the DNA looping of Series 1 and Series 2 constructs. As shown in the upper graph of Fig. 5A, the repression level observed in the presence of the sp TALE-Nhp6A (Fig. 5A, upper graph, green) is strongly altered relative to the data for the ns TALE-Nhp6A (Fig. 5A, upper graph, black). Interestingly, sp TALE-Nhp6A strongly assists repression looping at operator spacings near 132 bp and 143 bp (about one helical turn apart) but inhibits looping at operator spacings near 137 bp and 148 bp (about one helical turn apart). Maxima and minima are thus separated by ~ one half helical turn of DNA. The effect is emphasized when data are normalized to results in the presence of the ns TALE-Nhp6A protein (Fig. 5A, lower graph, green). These results clearly suggest that the DNA bend created by sp TALE-Nhp6A binding within the repression loop has an effect on repression that depends on the twist of the loop, with relaxed loops being further stabilized and twisted loops being further destabilized. This interpretation is again supported by thermodynamic model fitting (Table 1). The value of *K*_max_ for the sp TALE-Nhp6A (~64) is ~2-fold larger than in the presence of the ns TALE-Nhp6A or ns TALE. This confirms that sp TALE-Nhp6A enhances DNA looping at relaxed loops, overcoming the inhibitory effect of sp TALE binding by a striking 5.6-fold. Interestingly, while the Nhp6A domain acts by decreasing DNA resistance to bending, fit values of *C*_app_ (Table 1) suggest that sp TALE-Nhp6A does not overcome the twist inhibition caused by sp TALE binding to T_sp_ and may even pose a further obstacle (additional 50% *C*_app_ increase to 0.99 from 0.66).

**Fig 5.**
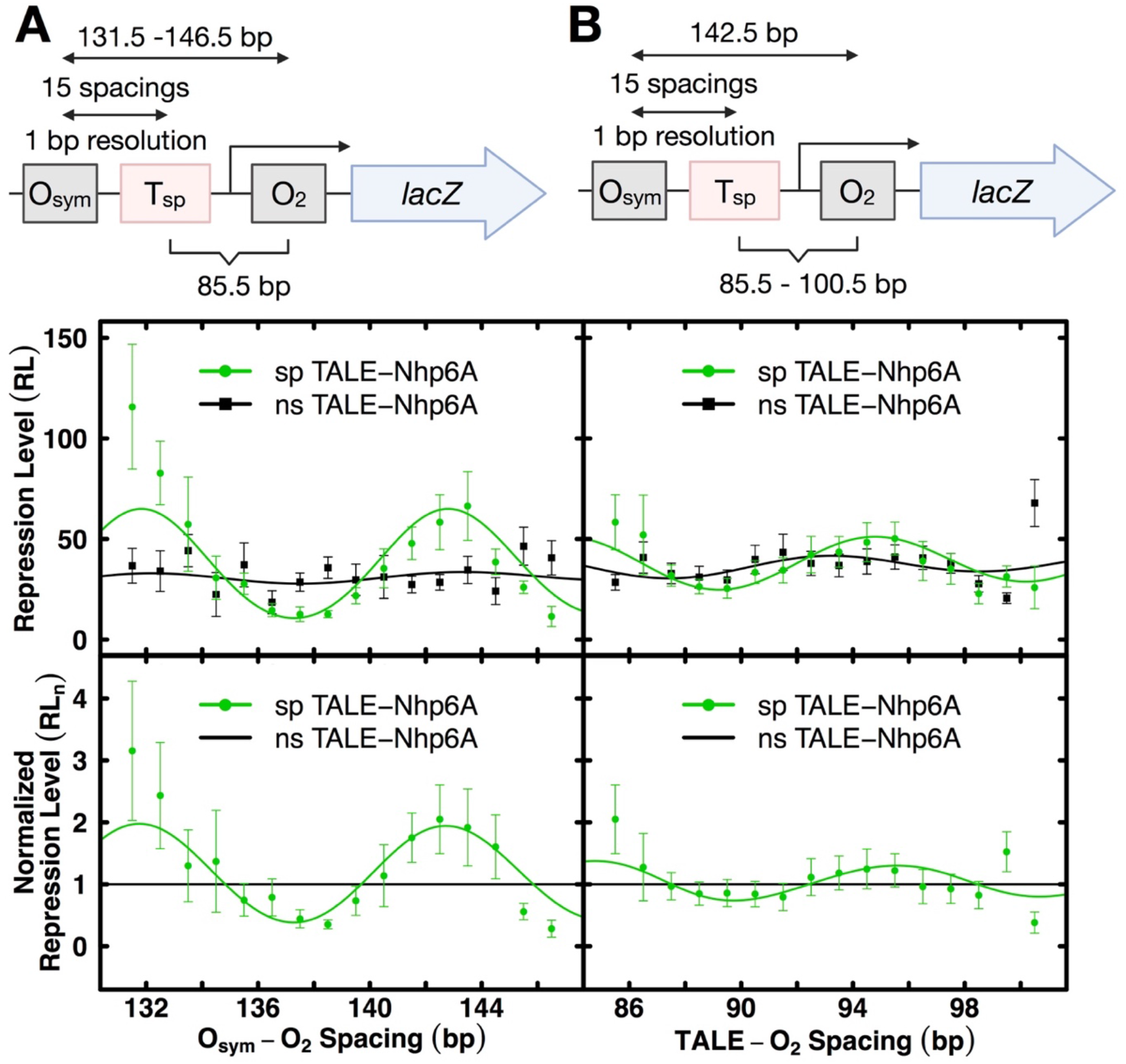
Effects of designed TALE-Nhp6A proteins on lacI-mediated DNA looping in living *E. coli*. A. Series 1 construct. The top diagram indicates control element configuration. The upper graph depicts repression level (*RL*) as a function of center-to-center lac operator spacing in the presence of the indicated nonspecific (black) or specific (red) TALE-Nhp6A protein where the TALE site is at a fixed position relative to O_2_ in the presence of lacI and 100 μM IPTG. The lower graph compares nonspecific or specific TALE-Nhp6A effects when repression level values are normalized to the corresponding values for nonspecific designed architectural protein (grey line indicates a value of 1.0). B. As in panel (A) except the data were obtained with series 2 constructs illustrated in top diagram. All depicted curve fits are to the thermodynamic model described in methods, with fit parameters reported in Table 1.

A phasing effect for TALE-Nhp6A binding is also observed in Series 2 reporter constructs where the TALE site is moved incrementally within a single untwisted repression loop (Fig. 5B, top diagram). Although phasing of the unoccupied TALE binding sequence itself changes loop stability (Fig. 5B, upper graph, black), the phasing effect is enhanced by sp TALE-Nhp6A binding. Optimal enhancement of the relaxed loop is observed at a T_sp_-O_2_ separation of ~95 bp. Thus, twist-dependent loop stabilization and destabilization are induced by sp TALE-Nhp6A binding within the repression loop, consistent with a sequence-targeted bending effect of sp TALE-Nhp6A (Supplemental Tables S5 and S6). Quantitative data fitting supports this interpretation (Table 1). ^TALE^*K*_Smax_ increases ~1.3 fold to 49.7 for sp TALE-Nhp6A (overall 3.2-fold change). ^TALE^*C*_app_increases from ~0.24 (ns TALE-Nhp6A) to ~0.36 (sp TALE-Nhp6A). This again indicates that TALE binding inhibits DNA twisting, an effect that Nhp6A fails to rescue. It should be noted that ^TALE^*C*_app_ values are lower in Series 2 than C_app_ in Series 1 (0.99) because the total loop size (142.5 bp) was deliberately chosen to be large (to accommodate multiple TALE positions) and near a looping probability maximum (relaxed loop).

### Modeling effects of artificial architectural proteins

To gain a better understanding of the observed effects of TALE and TALE-Nhp6A proteins on *lac* repression loop stabilization, we developed a series of molecular models based on a treatment of DNA at the level of base-pair steps with specific consideration of the spatial configuration of the bound repressor and designed architectural proteins (see Materials and Methods). We first generated energy-optimized configurations of repressor-mediated, TALE-free loops to identify the helical repeat in protein-free DNA that best mimics the chain-length dependent repression profiles observed in the absence of TALE binding. The looping propensities, i.e., *J* factors, extracted from the energies of chains with a 10.9-bp/turn helical repeat exhibit local maxima at center-to-center operator spacings of 133.5 and 144.5 bp and a local minimum at a spacing of 138.5 bp (Supplemental Fig. S4), values that match the repression measurements in Fig. 4A (series 1, grey data).

We next tested eight models of a TALE-bound construct derived from the high-resolution structure of the TAL effector PthXo1 in association with its DNA target sequence (11) to see which segments of the protein-DNA complex best capture the changes in amplitude and phasing of repression depicted in Fig. 4A (series 1, red data). While all of the models introduce a substantial reduction in the predicted ease of closing a DNA chain into a loop, only two exhibit a phase shift in the oscillatory variation of the *J* factor with operator spacing compared to TALE-free DNA (Supplemental Fig. S5). Moreover, the two representations of TALE-bound DNA yield looping profiles with maxima and minima matching the peaks and valleys in the repression data if the helical repeat of protein-free DNA is 10.8-bp/turn, with the change in intrinsic twist of protein-free DNA compensating for the ~11.5-bp/turn DNA repeat within the TALE assembly.

Finally we placed representative 10-bp fragments of Nhp6A-bound DNA on loops containing one of the selected TALE models. The fragments, taken from the collection of structures derived from NMR studies of the protein in complex with a 15-bp DNA duplex (25), were placed in two orientations and at variable locations downstream of the sp TALE construct on chains with a 10.7-10.8-bp helical repeat. The Nhp6A-induced bend and accompanying DNA undertwisting in the modeled pathways significantly enhance the looping propensities over those of TALE-free DNA in almost all cases (Supplemental Fig. S6). The sites of highest looping propensity depend upon the spacing between the TALE and Nhp6A binding sites, with maxima consistent with the observed peaks in repression (Fig. 5A, series 1, green data) occurring in constructs where the center of the Nhp6A-bound fragment lies 15 or 21 bp downstream of the center of the TALE-bound element and the protein-free segments have a 10.7-bp helical repeat. The enhancement pattern recurs but to a much lesser extent when Nhp6A is modeled an additional helical turn away, e.g., 25 bp from the TALE site. The looping propensities of the TALE-Nhp6A-bearing chains are also lower than those of TALE-less loops at certain spacings, e.g., in the vicinity of the minima when the Nhp6A lies 15 or 25 bp downstream of the TALE protein, in rough agreement with the repression data. The orientation of Nhp6A on DNA affects the site of protein uptake but does not change the character of the looping profiles. The predicted ease of looping is nearly identical for loops with Nhp6A bound in a reversed orientation 4 bp downstream of a site of forward binding (compare the profiles of loops with reversed vs. forward Nhp6A uptake — dashed vs. solid green lines — in Supplemental Fig. S6). Because the structure of the protein linker between the TALE and Nhp6A is unknown at high resolution, both Nhp6A dispositions, with their similar impacts, are considered.

Interestingly, the introduction of TALE-Nhp6A-bound fragments changes the predicted mix of configurational states adopted by the modeled DNA loops (Fig. 6, Supplemental Fig. S7). The loops formed most easily in the absence of architectural proteins, i.e., with 133.5 bp and 144.5 bp center-to-center operator spacing, follow antiparallel pathways. The DNA enters and exits the repressor in opposing directions, forming relatively smooth turns through apices located closer to one of the ends of the loops than the other (Fig. 6A). Whereas the loops labeled A1 depart the O_sym_ operator toward the center of the protein assembly — forming a turn closer to the 3′- than the 5′- end of the loop – the loops labeled A2 depart in the opposite direction, forming a turn closer to the 5′ than the 3′terminus of the loop. The specific TALE binding site thus is accommodated along the long, relatively straight stretch of the TALE-free A1 loops and at or near the turn in the TALE-free A2 loops. Introduction of a stiff, straight fragment of TALE-associated DNA at the designated binding site thus stabilizes the A1 compared to the A2 configuration, not only increasing the energy of the A2 form compared to the A1 form but also increasing the energy of the protein-free segments in the TALE-associated loops over those in the TALE-less loops. The increase in total energy gives rise to both the reduced looping propensities of the protein-bound vs. unbound loops and the difference in energy between the A1 and A2 configurations to the altered distribution of looped states (Fig. 6B). The specifically positioned TALE construct introduces more substantial distortions in the A2 than the A1 loops (see Supplemental Video S1).

**Fig. 6.**
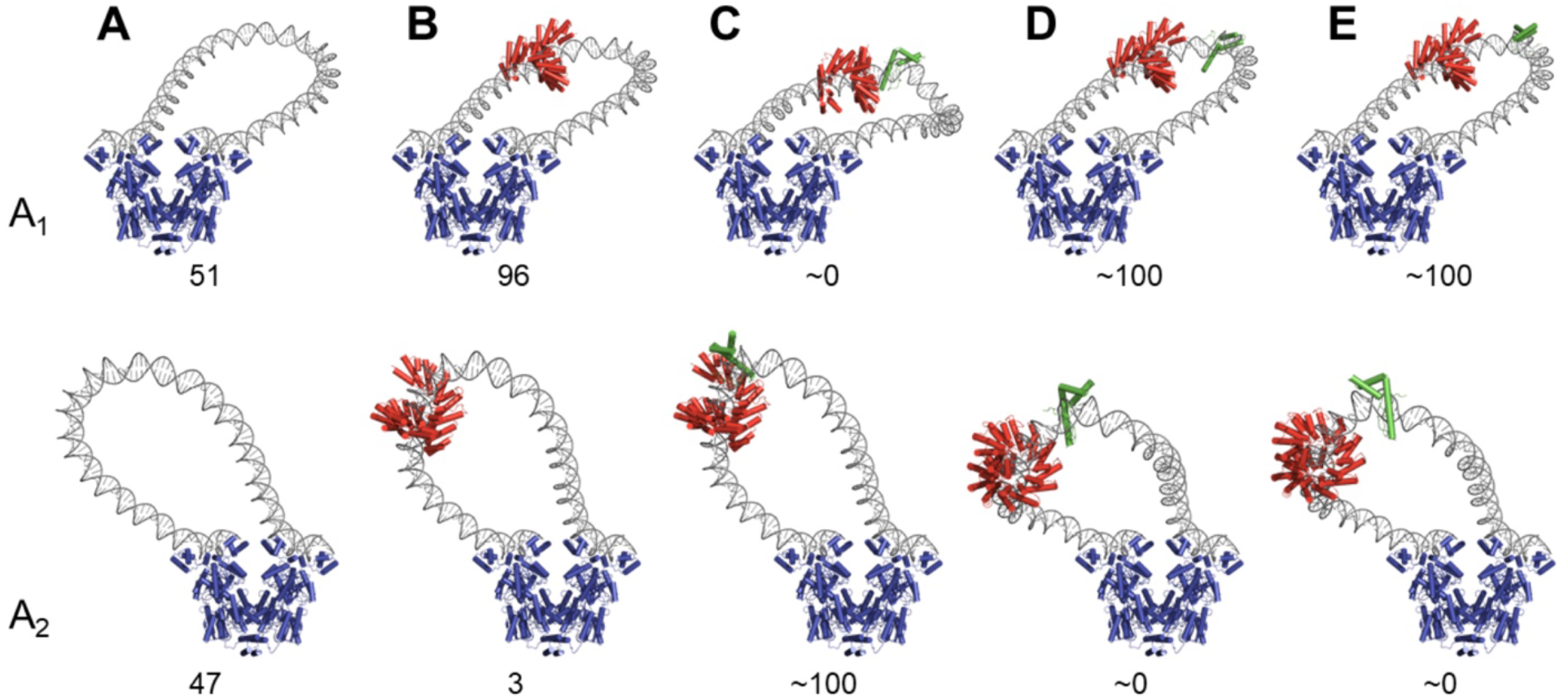
Models of DNA loop tuning by designed TALE-based architectural proteins. Energy-optimized configurations of Lac repressor-mediated DNA loops bearing TALE-Nhp6A constructs with O_sym_ and O_2_ operators spaced at distances of maximum looping propensity. The TALE construct is highlighted in red and the fused Nhp6A domain at predicted sites of likely binding in green. A. TALE-less antiparallel loops with 133.5-bp spacing and a 10.9-bp intrinsic helical repeat on protein-free DNA. B. TALE-associated loops with 131.5-bp spacing and a 10.8-bp repeat. C-E. TALE-Nhp6A constructs with 131.5-bp spacing, a 10.7-bp repeat, and Nhp6A spaced respectively 15, 21, and 25 bp downstream of the TALE protein. The Nhp6A in C and D associates with DNA in a forward orientation and that in E in a reverse orientation. The effects of protein binding on the predicted proportions of looped states are shown as percentage values below each image. Protein binding, however, has no effect on the structural family of these low-energy states. The pathway of the TALE-bound steps is represented by model 1 in Supplemental Table S7 and that of Nhp6A by model 4 from the ensemble of NMR-derived structures (pdb file 1j5n) (23). See Supplemental Table S9 for the *J* factors, structural families, and distances that must be spanned by the 65 amino acid residues separating the proline at the C terminus of the last repeat module of the TALE protein from the N-terminal methionine of Nhp6A in the dominant looped states.

The addition of Nhp6A to the TALE-bound models introduces further changes in the preferred looped states, stabilizing the A2 form significantly over the A1 form when Nhp6A is placed in a forward orientation 15 bp downstream of the TALE binding site (Fig. 6C) and enhancing the A1 form over the A2 form when placed in the same orientation 21 bp downstream (Fig. 6D). In other words, the placement and orientation of Nhp6A, depicted in green in Fig. 6, determine the balance between the looped states. Looping configurations predicted when Nhp6A is bound in a reversed orientation are similar to those found when the protein is placed 4 bp closer to the TALE protein in a forward orientation (Fig. 6E). TALE-Nhp6A binding is predicted to widen the mix of configurational states adopted by the least easily formed DNA loops. The protein-free loops are predicted to adopt a variety of high-energy pathways, including distorted forms of A1 and A2 looping from different structural families, as well as two loops, termed P1 and P2, with DNA entering and exiting the repressor in parallel directions. The P1 loop departs the O_sym_ operator in the same direction as the A1 loop, toward the central axis of the repressor, and the P2 loop in the same direction as the A2 loop, away from the repressor (see the variety of high-energy states in Fig. S7).

### Summary and implications

Beyond the intended goals of this study, our data, for the first time, demonstrate that TALE proteins strongly alter the bending and twisting properties of bound DNA, consistent with crowding of the DNA major groove by the TALE amino acids. Furthermore, our work gives insights into details of the mechanism of Nhp6A deformation of DNA. While tethered Nhp6A enhances DNA looping by inducing a targeted DNA kink, the direction of this deformation appears to be fixed and anisotropic, not creating a point of increased twist flexibility. These novel insights into the mechanical and dynamical properties of DNA bound by TALEs and/or Nhp6A would have been more difficult to observe and quantitate by other methods.

We believe that the combination of experimental gene repression data, thermodynamic modeling, and interpretation through biomechanical modeling of structure energetics, provides unusual insight into the behavior of *lac* repression loops modified by targeting of artificial architectural DNA binding proteins.

In summary, this work shows, for the first time, that sequence-specific DNA binding proteins based on TALEs and TALE fusions to the yeast Nhp6A protein can serve as targeted artificial architectural proteins that tune the probability of DNA looping in living bacteria by altering the mechanical properties of DNA.

## Funding

This work was supported by the Mayo Foundation, the Mayo Clinic Graduate School of Biomedical Sciences, and by National Institutes of Health grants GM34809 to WKO and GM75965 to LJM.

## Acknowledgments

We thank members of the Maher laboratory for assistance. We thank Marina Ramirez-Alvarado for sharing instrumentation.

## Cover Art

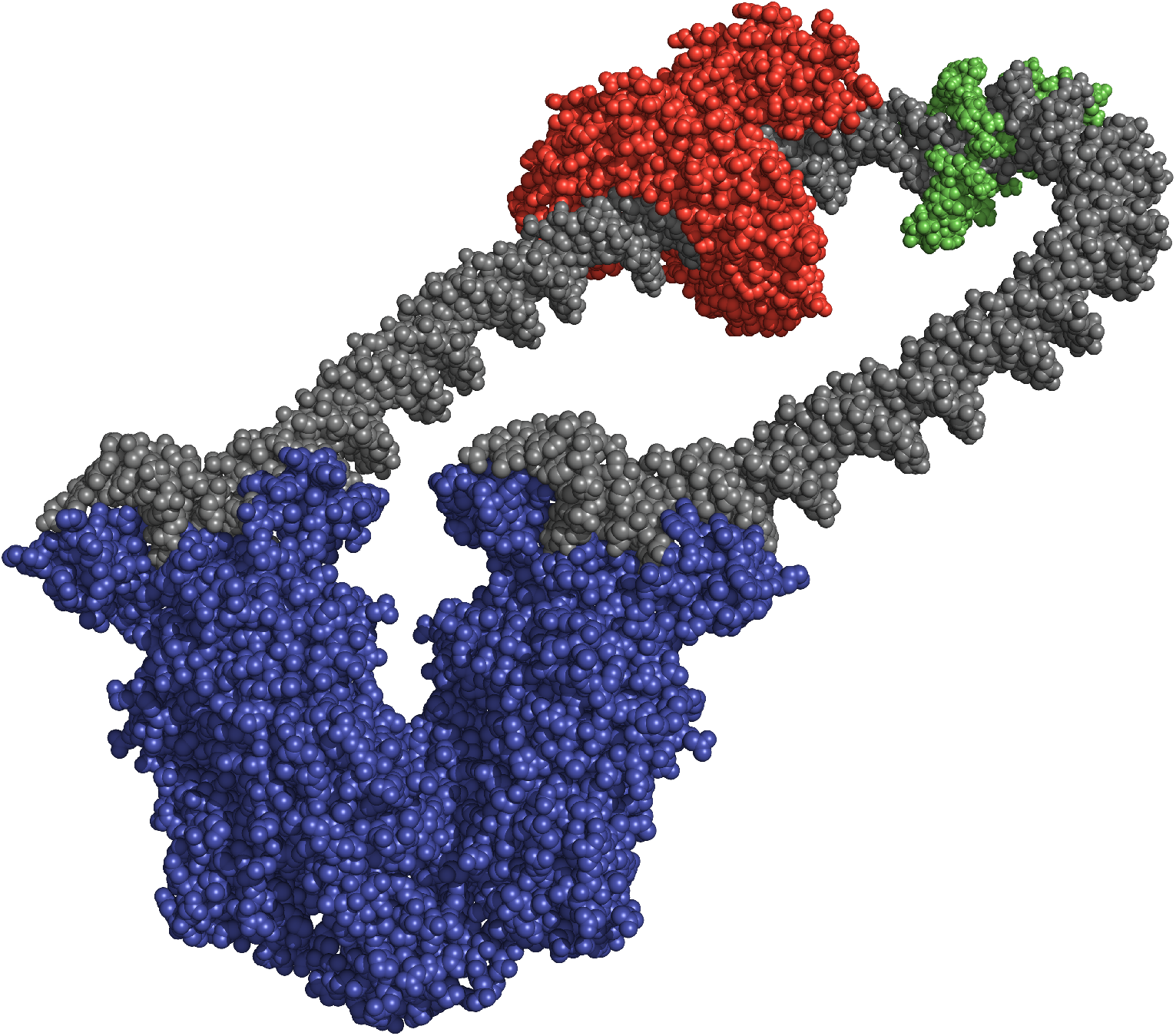
*Cover*: Molecular modeling based on biomechanical treatment of functional data from living *E. coli* bacteria for a 131.5-bp DNA repression loop anchored by the lactose repressor protein (blue). Tse et al. designed artificial architectural DNA binding proteins based on a sequence-specific Transcription Activator-Like Effector protein (red) with or without fusion to a sequence-nonspecific high mobility group DNA bending protein Nhp6A (green) to sculpt DNA physical properties and trajectory to promote or inhibit looping in cells. Rendering was accomplished using PyMol based on the configuration of protein-free DNA that optimizes the spatial arrangement of molecular fragments from PDB files 1efa, 1j5n, 1lbi, and 3ugm. Credit: R.T. Young. For more information see article by Tse, D. *et al.*, pages xxxx-xxxx in this issue.

## Notes

### Competing Interest Statement

The authors have declared no competing interest.

## References

1. Cantor, C.R. and Schimmel, P.R. (1980) Biophysical chemistry. W. H. Freeman, San Francisco.

2. Gelbart, W.M. and Knobler, C.M. (2009) Virology. Pressurized viruses. Science, 323, 1682–1683.

3. Hendrickson, W. and Schleif, R.F. (1984) Regulation of the Escherichia coli L-arabinose operon studied by gel electrophoresis DNA binding assay. J Mol Biol, 178, 611–628.

4. Jacob, F. and Monod, J. (1961) Genetic regulatory mechanisms in the synthesis of proteins. J Mol Biol, 3, 318–356.

5. Paull, T.T., Haykinson, M.J. and Johnson, R.C. (1993) The nonspecific DNA-binding and -bending proteins HMG1 and HMG2 promote the assembly of complex nucleoprotein structures. Genes Dev, 7, 1521–1534.

6. Zhou, G.L., Xin, L., Song, W., Di, L.J., Liu, G., Wu, X.S., Liu, D.P. and Liang, C.C. (2006) Active chromatin hub of the mouse alpha-globin locus forms in a transcription factory of clustered housekeeping genes. Mol Cell Biol, 26, 5096–5105.

7. Aki, T. and Adhya, S. (1997) Repressor induced site-specific binding of HU for transcriptional regulation. Embo J, 16, 3666–3674.

8. Becker, N.A. and Maher, L.J., 3rd. (2015) High-resolution mapping of architectural DNA binding protein facilitation of a DNA repression loop in Escherichia coli. Proc Natl Acad Sci U S A, 112, 7177–7182.

9. Boch, J. and Bonas, U. (2010) Xanthomonas AvrBs3 family-type III effectors: discovery and function. Annu Rev Phytopathol, 48, 419–436.

10. Moscou, M.J. and Bogdanove, A.J. (2009) A simple cipher governs DNA recognition by TAL effectors. Science, 326, 1501.

11. Mak, A.N., Bradley, P., Cernadas, R.A., Bogdanove, A.J. and Stoddard, B.L. (2012) The crystal structure of TAL effector PthXo1 bound to its DNA target. Science, 335, 716–719.

12. Rinaldi, F.C., Doyle, L.A., Stoddard, B.L. and Bogdanove, A.J. (2017) The effect of increasing numbers of repeats on TAL effector DNA binding specificity. Nucleic Acids Res, 45, 6960–6970.

13. Deng, D., Yan, C., Wu, J., Pan, X. and Yan, N. (2014) Revisiting the TALE repeat. Protein Cell, 5, 297–306.

14. Goodwin, G.H., Sanders, C. and Johns, E.W. (1973) A new group of chromatin-associated proteins with a high content of acidic and basic amino acids. Eur J Biochem, 38, 14–19.

15. Bustin, M. (2001) Revised nomenclature for high mobility group (HMG) chromosomal proteins. Trends Biochem Sci, 26, 152–153.

16. Reeves, R. (2010) Nuclear functions of the HMG proteins. Biochim Biophys Acta, 1799, 3–14.

17. Johns, E.W. (1982) The HMG chromosomal proteins. Academic Press, London; New York.

18. Bustin, M. and Reeves, R. (1996) High-mobility-group chromosomal proteins: architectural components that facilitate chromatin function. Prog Nucleic Acid Res Mol Biol, 54, 35–100.

19. Stojkova, P., Spidlova, P. and Stulik, J. (2019) Nucleoid-Associated Protein HU: A Lilliputian in Gene Regulation of Bacterial Virulence. Front Cell Infect Microbiol, 9, 159.

20. Lefebvre, V., Dumitriu, B., Penzo-Mendez, A., Han, Y. and Pallavi, B. (2007) Control of cell fate and differentiation by Sry-related high-mobility-group box (Sox) transcription factors. Int J Biochem Cell Biol, 39, 2195–2214.

21. Werner, M.H. and Burley, S.K. (1997) Architectural transcription factors: proteins that remodel DNA. Cell, 88, 733–736.

22. Thomas, J.O. and Travers, A.A. (2001) HMG1 and 2, and related ‘architectural’ DNA-binding proteins. Trends Biochem Sci, 26, 167–174.

23. Love, J.J., Li, X., Case, D.A., Giese, K., Grosschedl, R. and Wright, P.E. (1995) Structural basis for DNA bending by the architectural transcription factor LEF-1. Nature, 376, 791–795.

24. Czapla, L., Peters, J.P., Rueter, E.M., Olson, W.K. and Maher, L.J., 3rd. (2011) Understanding apparent DNA flexibility enhancement by HU and HMGB architectural proteins. J Mol Biol, 409, 278–289.

25. Masse, J.E., Wong, B., Yen, Y.M., Allain, F.H., Johnson, R.C. and Feigon, J. (2002) The S. cerevisiae architectural HMGB protein NHP6A complexed with DNA: DNA and protein conformational changes upon binding. J Mol Biol, 323, 263–284.

26. Hockemeyer, D., Wang, H., Kiani, S., Lai, C.S., Gao, Q., Cassady, J.P., Cost, G.J., Zhang, L., Santiago, Y., Miller, J.C. et al. (2011) Genetic engineering of human pluripotent cells using TALE nucleases. Nat Biotechnol, 29, 731–734.

27. Tesson, L., Usal, C., Menoret, S., Leung, E., Niles, B.J., Remy, S., Santiago, Y., Vincent, A.I., Meng, X., Zhang, L. et al. (2011) Knockout rats generated by embryo microinjection of TALENs. Nat Biotechnol, 29, 695–696.

28. Huang, P., Xiao, A., Zhou, M., Zhu, Z., Lin, S. and Zhang, B. (2011) Heritable gene targeting in zebrafish using customized TALENs. Nat Biotechnol, 29, 699–700.

29. Becker, N.A., Schwab, T.L., Clark, K.J. and Maher, L.J., 3rd. (2018) Bacterial gene control by DNA looping using engineered dimeric transcription activator like effector (TALE) proteins. Nucleic Acids Res, 46, 2690–2696.

30. Whipple, F.W. (1998) Genetic analysis of prokaryotic and eukaryotic DNA-binding proteins in Escherichia coli. Nucleic Acids Res, 26, 3700–3706.

31. Peters, J.P., Becker, N.A., Rueter, E.M., Bajzer, Z., Kahn, J.D. and Maher, L.J., 3rd. (2011) Quantitative methods for measuring DNA flexibility in vitro and in vivo. Methods Enzymol, 488, 287–335.

32. Becker, N.A., Kahn, J.D. and Maher, L.J., 3rd. (2005) Bacterial repression loops require enhanced DNA flexibility. J Mol Biol, 349, 716–730.

33. Bond, L.M., Peters, J.P., Becker, N.A., Kahn, J.D. and Maher, L.J., 3rd. (2010) Gene repression by minimal lac loops in vivo. Nucleic Acids Res, 38, 8072–8082.

34. Mossing, M.C. and Record, M.T., Jr. (1986) Upstream operators enhance repression of the lac promoter. Science, 233, 889–892.

35. Record, M.T., Jr., Mazur, S.J., Melancon, P., Roe, J.H., Shaner, S.L. and Unger, L. (1981) Double helical DNA: conformations, physical properties, and interactions with ligands. Annu Rev Biochem, 50, 997–1024.

36. Bellomy, G.R., Mossing, M.C. and Record, M.T., Jr. (1988) Physical properties of DNA in vivo as probed by the length dependence of the lac operator looping process. Biochemistry, 27, 3900–3906.

37. Becker, N.A., Kahn, J.D. and Maher, L.J., 3rd. (2007) Effects of nucleoid proteins on DNA repression loop formation in Escherichia coli. Nucleic Acids Res, 35, 3988–4000.

38. Becker, N.A., Kahn, J.D. and Maher, L.J., 3rd. (2008) Eukaryotic HMGB proteins as replacements for HU in E. coli repression loop formation. Nucleic Acids Res, 36, 4009–4021.

39. Becker, N.A., Greiner, A.M., Peters, J.P. and Maher, L.J., 3rd. (2014) Bacterial promoter repression by DNA looping without protein-protein binding competition. Nucleic Acids Res, 42, 5495–5504.

40. Clauvelin, N. and Olson, W.K. (2021) Synergy between Protein Positioning and DNA Elasticity: Energy Minimization of Protein-Decorated DNA Minicircles. J Phys Chem B, 125, 2277–2287.

41. Dickerson, R.E. (1989) Definitions and nomenclature of nucleic acid structure parameters. J Biomol Struct Dyn, 6, 627–634.

42. Lu, X.J. and Olson, W.K. (2003) 3DNA: a software package for the analysis, rebuilding and visualization of three-dimensional nucleic acid structures. Nucleic Acids Res, 31, 5108–5121.

43. Lu, X.J. and Olson, W.K. (2008) 3DNA: a versatile, integrated software system for the analysis, rebuilding and visualization of three-dimensional nucleic-acid structures. Nat Protoc, 3, 1213–1227.

44. Lewis, M., Chang, G., Horton, N.C., Kercher, M.A., Pace, H.C., Schumacher, M.A., Brennan, R.G. and Lu, P. (1996) Crystal structure of the lactose operon repressor and its complexes with DNA and inducer. Science, 271, 1247–1254.

45. Spronk, C.A., Bonvin, A.M., Radha, P.K., Melacini, G., Boelens, R. and Kaptein, R. (1999) The solution structure of Lac repressor headpiece 62 complexed to a symmetrical lac operator. Structure, 7, 1483–1492.

46. Bell, C.E. and Lewis, M. (2000) A closer view of the conformation of the Lac repressor bound to operator. Nat Struct Biol, 7, 209–214.

47. Romanuka, J., Folkers, G.E., Biris, N., Tishchenko, E., Wienk, H., Bonvin, A.M., Kaptein, R. and Boelens, R. (2009) Specificity and affinity of Lac repressor for the auxiliary operators O2 and O3 are explained by the structures of their protein-DNA complexes. J Mol Biol, 390, 478–489.

48. Czapla, L., Swigon, D. and Olson, W.K. (2006) Sequence-Dependent Effects in the Cyclization of Short DNA. J Chem Theory Comput, 2, 685–695.

49. Du, Q., Smith, C., Shiffeldrim, N., Vologodskaia, M. and Vologodskii, A. (2005) Cyclization of short DNA fragments and bending fluctuations of the double helix. Proc Natl Acad Sci U S A, 102, 5397–5402.

50. Horowitz, D.S. and Wang, J.C. (1984) Torsional rigidity of DNA and length dependence of the free energy of DNA supercoiling. J Mol Biol, 173, 75–91.

51. Heath, P.J., Clendenning, J.B., Fujimoto, B.S. and Schurr, J.M. (1996) Effect of bending strain on the torsion elastic constant of DNA. J Mol Biol, 260, 718–730.

52. Geanacopoulos, M., Vasmatzis, G., Zhurkin, V.B. and Adhya, S. (2001) Gal repressosome contains an antiparallel DNA loop. Nat Struct Biol, 8, 432–436.

53. Colasanti, A.V., Grosner, M.A., Perez, P.J., Clauvelin, N., Lu, X.J. and Olson, W.K. (2013) Weak operator binding enhances simulated Lac repressor-mediated DNA looping. Biopolymers, 99, 1070–1081.

54. Perez, P.J. and Olson, W.K. (2016) Insights into Genome Architecture Deduced from the Properties of Short Lac Repressor-mediated DNA Loops. Biophys Rev, 8, 135–144.

55. Perez, P.J., Clauvelin, N., Grosner, M.A., Colasanti, A.V. and Olson, W.K. (2014) What controls DNA looping? Int J Mol Sci, 15, 15090–15108.

