## Supplementary data for "Designed architectural proteins that tune DNA looping in bacteria"

Tanya L. Schwab[1], Karl J. Clark[1], and L. James Maher, III[1]*

[1]Department of Biochemistry and Molecular Biology
Mayo Clinic College of Medicine and Science
200 First St. SW
Rochester, MN 55905, USA

[2]Department of Chemistry and Chemical Biology
Rutgers, the State University of New Jersey
Center for Quantitative Biology
Piscataway, NJ 08854, USA

[3]Department of Chemistry and Biochemistry
University of Northern Iowa
1227 West 27th Street
Cedar Falls, IA 50614-0423

Supplemental Tables S5 & S6 are available within a separate excel file.
Supplemental Video S1 is available as a separate .mp4 file.

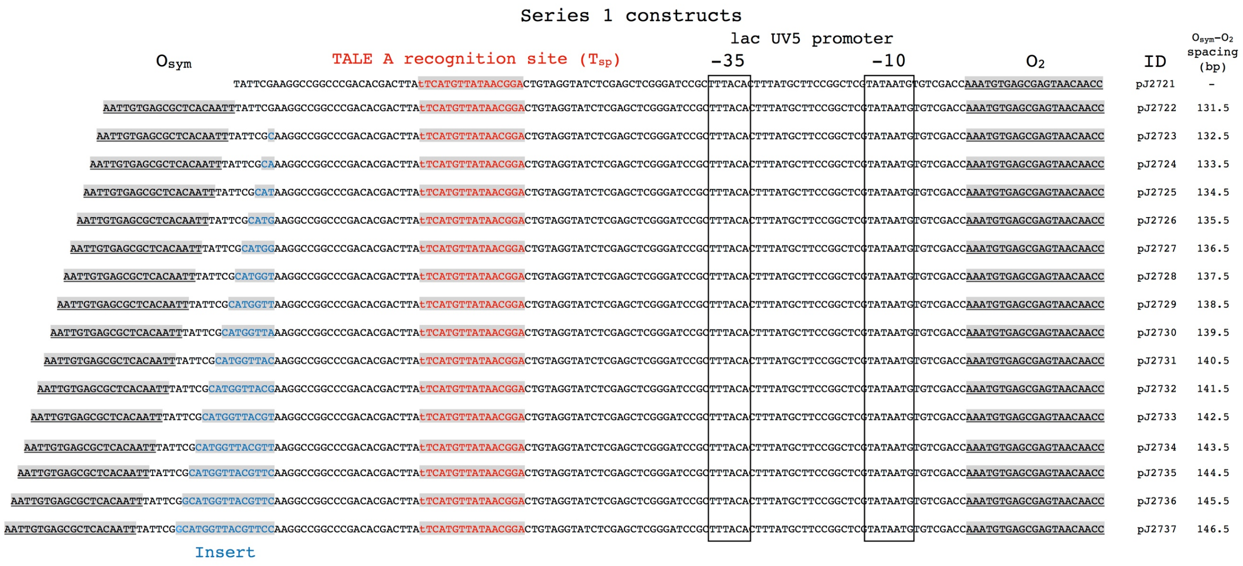

**Supplemental Fig. S1.** Sequences of control region elements of Series 1 reporter constructs. Key elements are shaded grey or boxed, as indicated. TALE binding sites, $T_{sp}$, are shown in red. Thymine residue recognized by the initial TALE invariant module is indicated in lower case. Operator spacings (center-to-center) are indicated for the shaded and underlined sequences.

#### Series 2 constructs

Labels above the sequences: $O_{sym}$ · TALE A recognition site ($T_{sp}$) · lac UV5 promoter (−35, −10) · $O_2$

| Sequence | ID | $T_{sp}$–$O_2$ spacing (bp) |
|---|---|---|
| AATTGTGAGCGCTCACAATTTATTCGGCATGGTCCTAGGGGCCGGCCCGACACGACTTAtTCATGTTATAACGGACTGTAGGTATCTCGAGCTCGGGATCCGCTTTACACTTTATGCTTCCGGCTCGTATAATGTGTCGACCAAATGTGAGCGAGTAACAACC | pJ2748 | 85.5 |
| AATTGTGAGCGCTCACAATTTATTCGGCATGGTCCTAGGGGCCGCCCGACACGACTTAtTCATGTTATAACGGAGCTGTAGGTATCTCGAGCTCGGGATCCGCTTTACACTTTATGCTTCCGGCTCGTATAATGTGTCGACCAAATGTGAGCGAGTAACAACC | pJ2749 | 86.5 |
| AATTGTGAGCGCTCACAATTTATTCGGCATGGTCCTAGGGGCCGCCGACACGACTTAtTCATGTTATAACGGAGCCTGTAGGTATCTCGAGCTCGGGATCCGCTTTACACTTTATGCTTCCGGCTCGTATAATGTGTCGACCAAATGTGAGCGAGTAACAACC | pJ2750 | 87.5 |
| AATTGTGAGCGCTCACAATTTATTCGGCATGGTCCTAGGGGCCGCGACACGACTTAtTCATGTTATAACGGAGCCCTGTAGGTATCTCGAGCTCGGGATCCGCTTTACACTTTATGCTTCCGGCTCGTATAATGTGTCGACCAAATGTGAGCGAGTAACAACC | pJ2751 | 88.5 |
| AATTGTGAGCGCTCACAATTTATTCGGCATGGTCCTAGGGGCCGGACACGACTTAtTCATGTTATAACGGAGCCCCTGTAGGTATCTCGAGCTCGGGATCCGCTTTACACTTTATGCTTCCGGCTCGTATAATGTGTCGACCAAATGTGAGCGAGTAACAACC | pJ2752 | 89.5 |
| AATTGTGAGCGCTCACAATTTATTCGGCATGGTCCTAGGGGCCGACACGACTTAtTCATGTTATAACGGAGCCCGCTGTAGGTATCTCGAGCTCGGGATCCGCTTTACACTTTATGCTTCCGGCTCGTATAATGTGTCGACCAAATGTGAGCGAGTAACAACC | pJ2753 | 90.5 |
| AATTGTGAGCGCTCACAATTTATTCGGCATGGTCCTAGGGGCCGCACGACTTAtTCATGTTATAACGGAGCCCGACTGTAGGTATCTCGAGCTCGGGATCCGCTTTACACTTTATGCTTCCGGCTCGTATAATGTGTCGACCAAATGTGAGCGAGTAACAACC | pJ2754 | 91.5 |
| AATTGTGAGCGCTCACAATTTATTCGGCATGGTCCTAGGGGCCGACGACTTAtTCATGTTATAACGGAGCCCGACCTGTAGGTATCTCGAGCTCGGGATCCGCTTTACACTTTATGCTTCCGGCTCGTATAATGTGTCGACCAAATGTGAGCGAGTAACAACC | pJ2755 | 92.5 |
| AATTGTGAGCGCTCACAATTTATTCGGCATGGTCCTAGGGGCCGCGACTTAtTCATGTTATAACGGAGCCCGACACTGTAGGTATCTCGAGCTCGGGATCCGCTTTACACTTTATGCTTCCGGCTCGTATAATGTGTCGACCAAATGTGAGCGAGTAACAACC | pJ2756 | 93.5 |
| AATTGTGAGCGCTCACAATTTATTCGGCATGGTCCTAGGGGCCGGACTTAtTCATGTTATAACGGAGCCCGACACCTGTAGGTATCTCGAGCTCGGGATCCGCTTTACACTTTATGCTTCCGGCTCGTATAATGTGTCGACCAAATGTGAGCGAGTAACAACC | pJ2757 | 94.5 |
| AATTGTGAGCGCTCACAATTTATTCGGCATGGTCCTAGGGGCCGACTTAtTCATGTTATAACGGAGCCCGACACGCTGTAGGTATCTCGAGCTCGGGATCCGCTTTACACTTTATGCTTCCGGCTCGTATAATGTGTCGACCAAATGTGAGCGAGTAACAACC | pJ2758 | 95.5 |
| AATTGTGAGCGCTCACAATTTATTCGGCATGGTCCTAGGGGCCGCTTAtTCATGTTATAACGGAGCCCGACACGACTGTAGGTATCTCGAGCTCGGGATCCGCTTTACACTTTATGCTTCCGGCTCGTATAATGTGTCGACCAAATGTGAGCGAGTAACAACC | pJ2759 | 96.5 |
| AATTGTGAGCGCTCACAATTTATTCGGCATGGTCCTAGGGGCCGTTAtTCATGTTATAACGGAGCCCGACACGACCTGTAGGTATCTCGAGCTCGGGATCCGCTTTACACTTTATGCTTCCGGCTCGTATAATGTGTCGACCAAATGTGAGCGAGTAACAACC | pJ2760 | 97.5 |
| AATTGTGAGCGCTCACAATTTATTCGGCATGGTCCTAGGGGCCGTAtTCATGTTATAACGGAGCCCGACACGACTCTGTAGGTATCTCGAGCTCGGGATCCGCTTTACACTTTATGCTTCCGGCTCGTATAATGTGTCGACCAAATGTGAGCGAGTAACAACC | pJ2761 | 98.5 |
| AATTGTGAGCGCTCACAATTTATTCGGCATGGTCCTAGGGGCCGAtTCATGTTATAACGGAGCCCGACACGACTTCTGTAGGTATCTCGAGCTCGGGATCCGCTTTACACTTTATGCTTCCGGCTCGTATAATGTGTCGACCAAATGTGAGCGAGTAACAACC | pJ2762 | 99.5 |
| AATTGTGAGCGCTCACAATTTATTCGGCATGGTCCTAGGGGCCGtTCATGTTATAACGGAGCCCGACACGACTTACTGTAGGTATCTCGAGCTCGGGATCCGCTTTACACTTTATGCTTCCGGCTCGTATAATGTGTCGACCAAATGTGAGCGAGTAACAACC | pJ2763 | 100.5 |

**Supplemental Fig. S2**. Sequences of control region elements of Series 2 reporter constructs. Key elements are shaded grey or boxed, as indicated. TALE binding sites, $T_{sp}$, are shown in red. Thymine residue recognized by the initial TALE invariant module is indicated in lower case. $O_2$ - TALE spacings (center-to-center) are indicated for the shaded and underlined sequences.

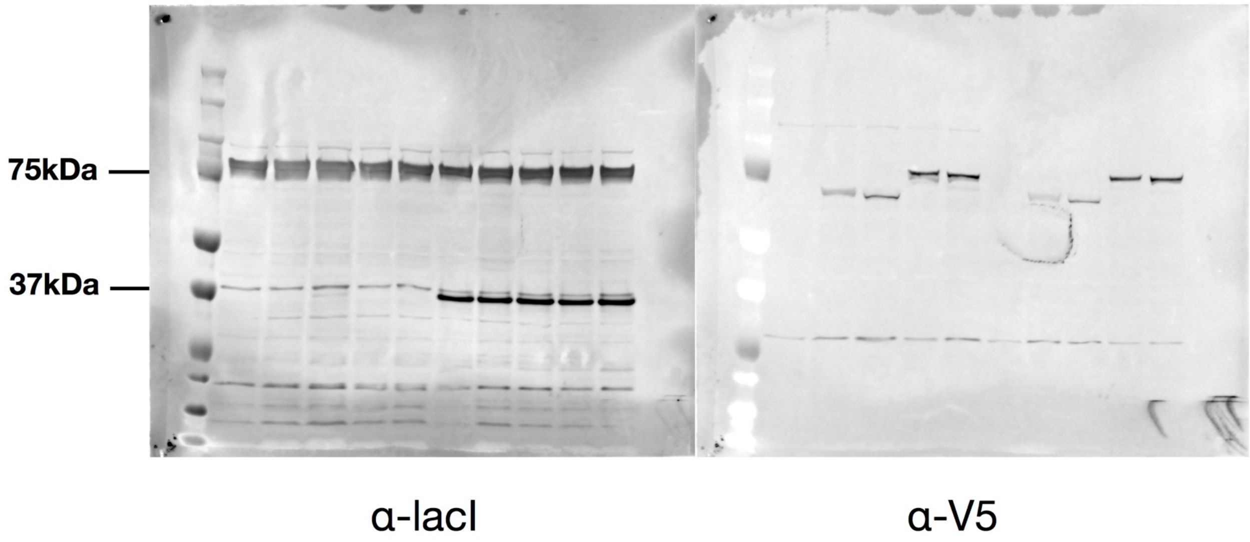

**Supplemental Fig. S3**. Uncropped images related to Fig. 3A.

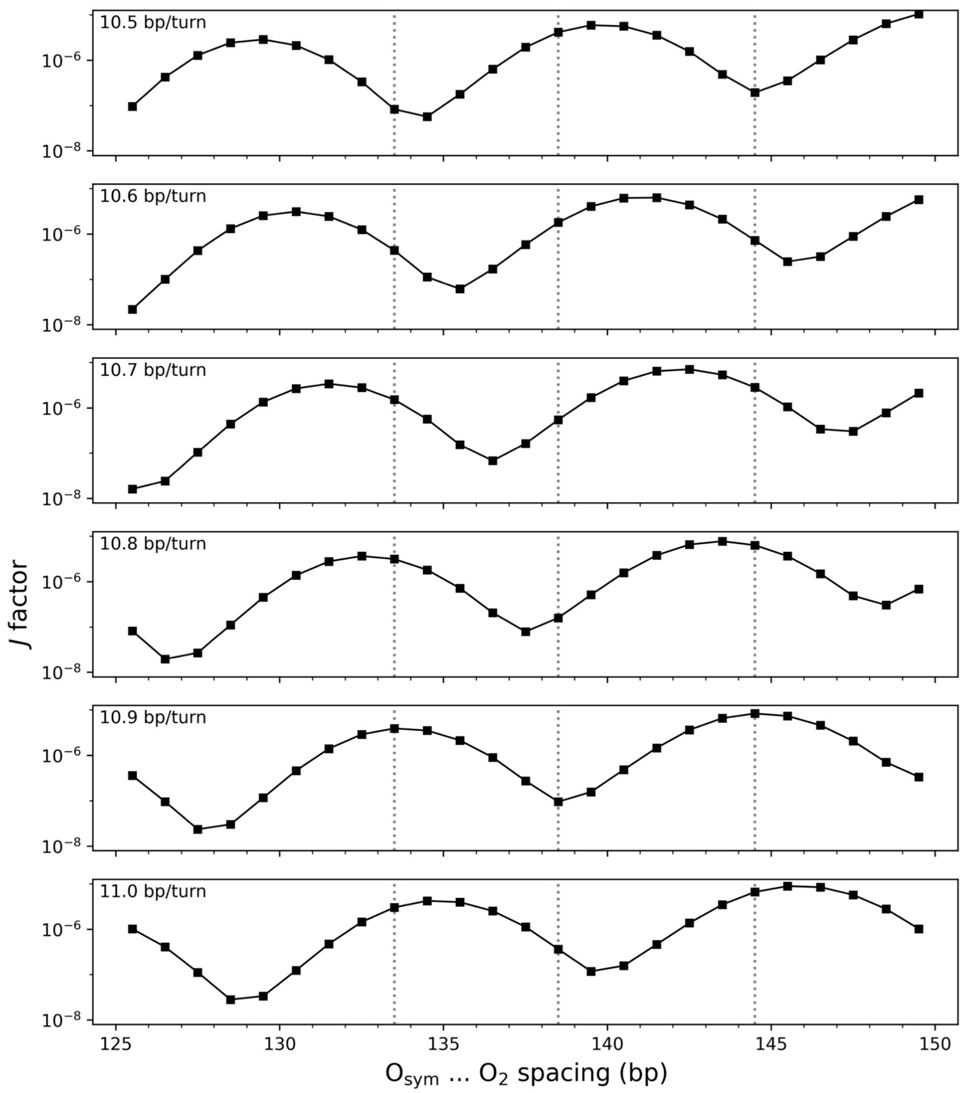

**Supplemental Fig. S4**. A 10.9-bp/turn DNA helical repeat yields looping profiles consistent with the chain-length dependent repression levels measured for TALE-free DNA. *J* factors of Lac repressor-mediated loops extracted from the energies of optimized loop configurations as a function of operator spacing. Protein-free DNA treated as an ideal, inextensible, naturally straight chain with elastic properties characteristic of mixed-sequence DNA and a 10.5 to 11 bp/turn helical repeat in its equilibrium rest state. The spacings associated with local maxima/minima in repression of TALE-less DNA (Fig. 4A: series I, grey data) are highlighted by dotted grey lines.

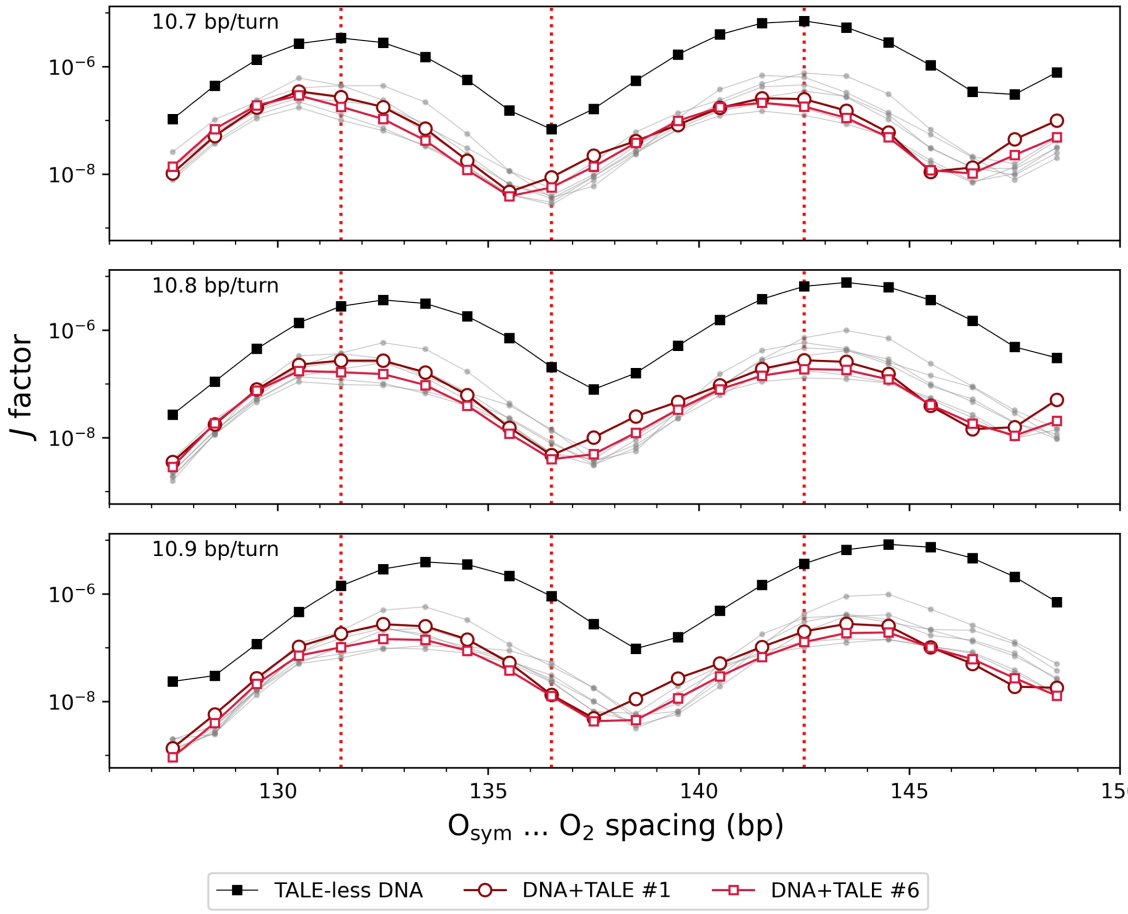

**Supplemental Fig. S5**. Selected 16-bp fragments from the complex of DNA with the TALE PthXo1 (1) introduce changes in DNA looping that mirror the changes in the amplitude and phasing of repression found in TALE-bound DNA. *J* factors of Lac repressor-mediated loops bearing TALE-bound elements extracted from the cited high-resolution structure and placed 85.5 bp upstream of the O2 operator in chains with 126.5-148.5 bp operator spacings and protein-free DNA with a 10.7-10.9-bp/turn helical repeat. See Supplemental Table S7 for the components of the high-resolution TALE-DNA complex incorporated in the modeled fragments. The looping profiles associated with the models 1 and 6, depicted respectively as large red circles and small red squares, show both a reduction in amplitude and a phase shift of the type shown in the repression levels of TALE-bound vs. TALE-less DNA. The models illustrated in light gray do not show a phase shift compared to TALE-less DNA (black squares). The spacing associated with local maxima/minima in repression of TALE-bound DNA (Fig. 4A: series I, red data) are highlighted by dotted red lines.

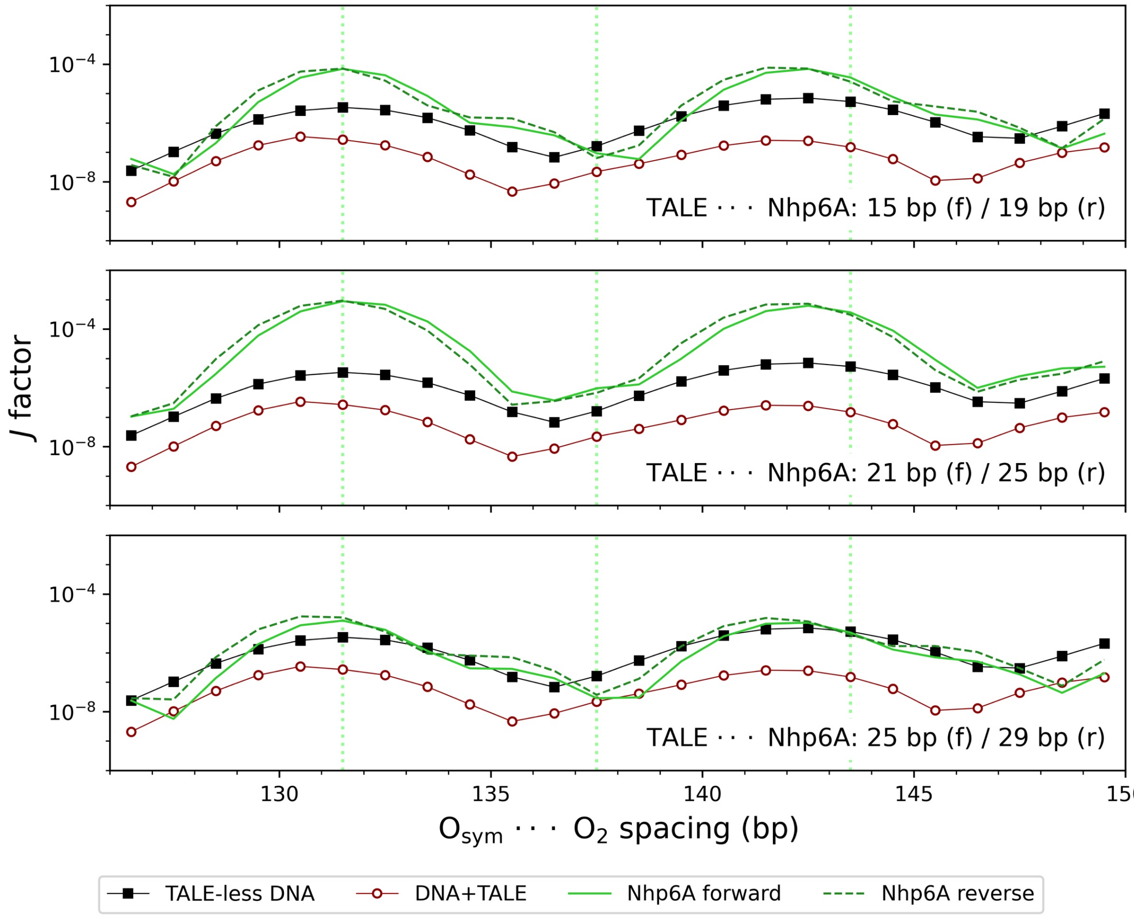

**Supplemental Fig. S6**. Selected settings of Nhp6A-bound DNA relative to the TALE construct on DNA introduce extremes of looping enhancement/depression of the type seen in the chain-length-dependent repression of loops bearing TALE-Nhp6A constructs. *J* factors of Lac repressor-mediated loops bearing TALE-Nhp6A constructs upstream of the O2 operator in chains with 126.5-148.5-bp operator spacings and protein-free DNA with a 10.7 bp helical repeat. The Nhp6A lies downstream of the TALE construct in forward and reverse settings (profiles depicted by solid/dashed green lines). The spacing between proteins is expressed in terms of the number of base pairs between the centers of TALE and Nhp6A uptake on DNA. The profiles of TALE-less (black squares) and TALE-bound (red circles) DNA are shown for comparison. The spacings associated with local maxima/minima in repression of TALE-Nhp6A-bound DNA (Fig. 5A: series I, green data) are highlighted by dotted green lines. Loops bearing TALE-bound elements described by model 6 are similar to those reported here using model 1 (see Supplemental Fig. S5). Chains modeled with a 10.8 bp helical repeat of protein-free DNA (data not shown) exhibit similar, albeit slightly out-of-phase, oscillatory looping profiles.

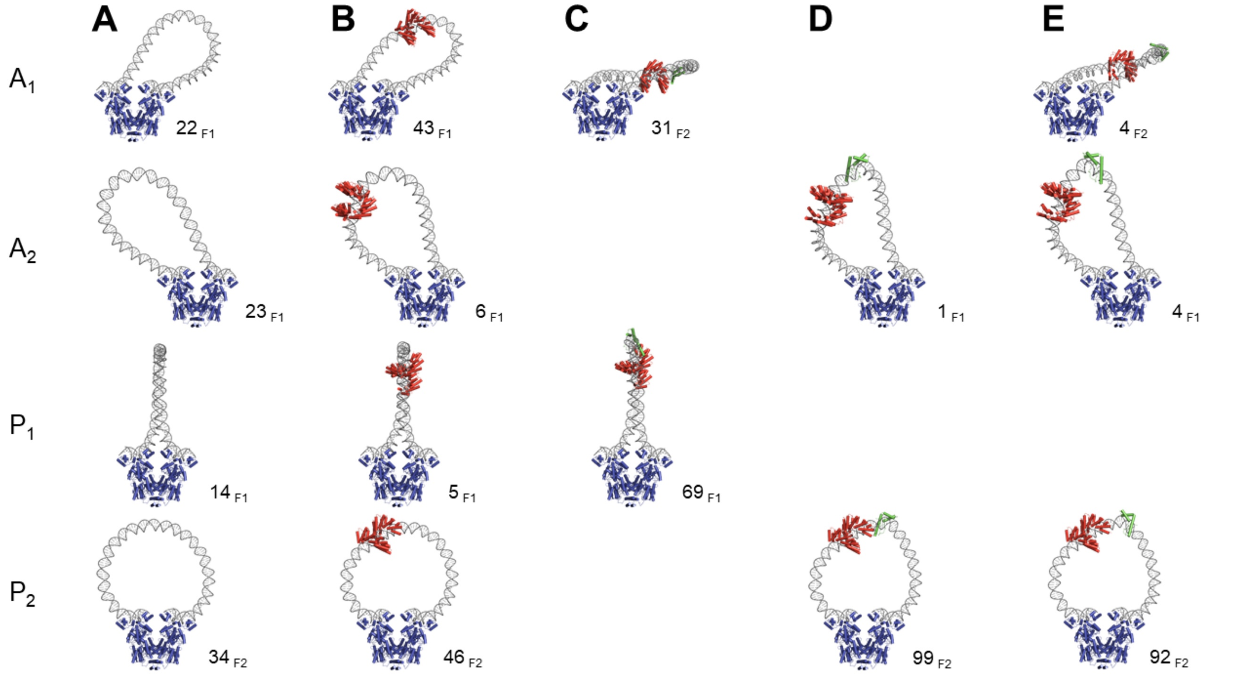

**Supplemental Fig. S7**. Designed TALE-based architectural proteins change the mix of configurational states found at points of minimum loop-closure propensity. The TALE construct is depicted in red and the fused Nhp6A domain in green. A. TALE-less antiparallel loops with 138.5-bp spacing and a 10.9-bp intrinsic helical repeat on protein-free DNA; B. TALE-associated loops with 136.5-bp spacing and a 10.8-bp repeat; C-E. TALE-Nhp6A constructs with 137.5-bp spacing, a 10.7-bp repeat, and Nhp6A incorporated respectively at sites 15, 21, and 25 bp downstream of the TALE protein. The Nhp6A in C and D associates with DNA in a forward orientation and that in E in a reverse orientation. Note the effects of protein uptake on the predicted proportions and configurational families of looped states (percentage values and footnoted nomenclature to the right of each image). The pathway of the TALE-bound steps is represented by model 1 in Supplemental Table S7 and that of Nhp6A by model 4 from the ensemble of NMR-derived structures (pdb file 1j5n) (2). See Supplemental Table S9 for the *J* factors and distances that must be spanned by the 65 amino acid residues separating the C terminus of proline in the last repeat module of the TALE from the N-terminal methionine of Nhp6A in the dominant looped states.

**Supplemental Video S1**.

Models of DNA loop tuning by designed TALE-based architectural proteins from different perspectives. This video, found in the accompanying mp4 file, highlights the configurational differences among Lac repressor-mediated loops bearing different TALE-Nhp6A constructs. The images convert from the conventional description of loop formation with respect to the V-shaped repressor assembly to different rotated viewpoints. Loops in $A_1$ and $A_2$ configurations progress from TALE-less to TALE-bound and TALE-Nh6pA-bound structures with 15, 21, and 25 bp between the centers of the TALE- and Nhp6A-bound sites. The table below gives the times when optimized structures with the specified molecular components appear in the video. See the legend to Fig. 6 for further details.

| Video time index (sec) | DNA | TALE | Nhp6A 15 bp forward | Nhp6A 21 bp forward | Nhp6A 25 bp reverse |
|---|---|---|---|---|---|
| Conventional view showing V-shaped Lac repressor | | | | | |
| 03:00 | + | | | | |
| 04:00 | + | + | | | |
| 05:00 | + | + | + | | |
| 06:00 | + | + | | + | |
| 08:00 | + | + | | | + |
| View in the direction of the operator axes following rotation about the vertical axis | | | | | |
| 10:00 | + | | | | |
| 12:00 | + | + | | | |
| 13:00 | + | + | + | | |
| 15:00 | + | + | | + | |
| 16:00 | + | + | | | + |
| View into the V of the repressor following rotations about the horizontal and normal axes | | | | | |
| 21:00 | + | | | | |
| 23:00 | + | + | | | |
| 25:00 | + | + | + | | |
| 26:00 | + | + | | + | |
| 27:00 | + | + | | | + |

SUPPORTING INFORMATION METHODS

### Treatment of protein-bound DNA

The Lac repressor assembly is represented by a rigid, V-shaped model, constructed by superposition of the structures of two well-resolved macromolecular complexes (3,4) —the 2.6-Å resolution X-ray crystallographic structure of the Lac protein dimer bound to the $O_{sym}$ operator (pPDB: 1efa) (5) and the 2.7-Å resolution X-ray structure of the tetrameric form of the protein without DNA-binding headpieces (PDB: 1lbi) (6). The DNA operators are assigned the rigid-body parameters adopted in the former complex, an approximation based on the similar spatial arrangements and comparable numbers of close protein-DNA contacts identified in NMR solution studies of the $O_{sym}$ and $O_2$ operators bound to the N-terminal headpieces of the repressor protein (PDB: 1cjg and 2kej, respectively) (7,8). The overall deformation of the 15-bp modeled operators (see below), however, is somewhat greater than that deduced from the NMR data, introducing a greater bend between terminal base pairs compared to the corresponding stretches of $O_{sym}$ and $O_2$ in the structural ensemble (70° in the current model vs. respective mean values of $41_{\pm 10}$° and $50_{\pm 6}$° in the NMR structures) but overwinding rather than underwinding (+28.9° vs. –14.7° and –22.6° over the central 14-bp steps) relative to the same length of ideal B DNA. Here the uptake of twist is measured in terms of the twist of supercoiling (9,10), with the ideal chain assigned a twist of 360°/10.5 = 34.3° per base-pair step.

The model of the TALE construct is based on the configurations of protein-DNA fragments found within the 3.0-Å structure of the TALE PthXo1 from the rice pathogen *Xanthomonas oryzae* (PDB: 3ugm) in association with its naturally occurring DNA target sequence (1). The 36-bp duplex is broken into eight overlapping 16-bp fragments, each found using the SNAP component of the 3DNA/DSSR software (11,12) to have base-pair atoms in hydrogen-bonded contact with amino-acid atoms on the TAL effector (Supplemental Table S7) and each assigned the set of rigid-body parameters extracted from the corresponding DNA pathway. The Nhp6A-bound element is taken from the ensemble of 20 structures deduced from NMR studies of the complex of protein with a 15-bp DNA duplex containing a recognition sequence for the HMGB transcription factor SRY (PDB: 1j5n) (2). The protein-bound DNA is shortened to the ten base pairs found with SNAP (11,12) to be in direct hydrogen-bonded contact with atoms on protein, i.e., omitting the three base pairs at the 5′-ends and the two base pairs at the 3′-ends of the deposited structures. The Nhp6A-bound fragment is described, for simplicity, by two representative pathways (models 4 and 9 within the pdb file) with 47 and 48 of the 54 rigid-body parameters within one standard deviation of the average values at each step in the structural ensemble (Supplemental Table S8). Without prior knowledge of how and where the protein associates with the DNA loop, the Nhp6A-bound fragment is placed in two orientations at different locations with respect to the TALE-bound fragment. The protein is introduced so that the positively charged N-terminal tail of Nhp6A makes initial contact to DNA via either the leading strand or the complementary strand, here respectively termed forward vs. reverse orientation, with up to 28 bp separating the center of the Nhp6A-bound site from that of the TALE target site. Like the Lac repressor and the TAL effector, Nhp6A is treated as a ‘side group’ of DNA with the coordinates of amino-acid atoms expressed in the reference frame of one of the associated base pairs. Whereas the repressor slightly overwinds DNA, the TALE and Nhp6A proteins underwind DNA, with respective helical repeats of 10.4, 11.7, and 12.8 bp/turn (values obtained from the quotients of 360° and the average twists of supercoiling — 34.7°, 30.7°, and 28.2°).

The operators are aligned against the repressor such that the large protein-induced kink of DNA within each bound fragment lies on the CG base-pair step found to take up the distortion in the NMR structural ensembles (7,8). That is, whereas the central base-pair step of the modeled $O_{sym}$ operator coincides with the highly kinked CG step at the center of the experimentally characterized system, the kink in the modeled $O_2$ operator lies off-center, on the CG step that incorporates the central G of the leading strand. The protein-bound DNA is restricted to the 15-bp steps in direct contact with repressor, with the three base pairs at the 3′-end of $O_{sym}$ and the two base pairs at the 5′-end of $O_2$ treated as protein-free DNA. The three base pairs at the 5′-end of $O_{sym}$ and the four base pairs at the 3′-end of $O_2$ are not included in the model.

**Supplemental Table S1**. TALE recognition sequences

| TALE | recognition sequence |
|---|---|
| A (sp) | tTCATGTTATAACGGA |
| C (ns) | tTACAAGTGGCTCATT |

**Supplemental Table S2**. Series 1 spacing construct plasmids used in this study

| Plasmid | Distal Operator | Spacing (bp)* | | | Proximal Operator | $T_{sp}$ |
|---|---|---|---|---|---|---|
| | | $O_{sym}\cdots O_2$ | $O_{sym}\cdots T_{sp}$ | $T_{sp}\cdots O_2$ | | |
| pJ2644 | - | - | - | - | $T_A$ | - |
| pJ2646 | - | - | - | - | $T_C$ | - |
| pJ2721 | - | - | - | 85.5 | $O_2$ | + |
| pJ2722 | $O_{sym}$ | 131.5 | 46 | 85.5 | $O_2$ | + |
| pJ2723 | $O_{sym}$ | 132.5 | 47 | 85.5 | $O_2$ | + |
| pJ2724 | $O_{sym}$ | 133.5 | 48 | 85.5 | $O_2$ | + |
| pJ2725 | $O_{sym}$ | 134.5 | 49 | 85.5 | $O_2$ | + |
| pJ2726 | $O_{sym}$ | 135.5 | 50 | 85.5 | $O_2$ | + |
| pJ2727 | $O_{sym}$ | 136.5 | 51 | 85.5 | $O_2$ | + |
| pJ2728 | $O_{sym}$ | 137.5 | 52 | 85.5 | $O_2$ | + |
| pJ2729 | $O_{sym}$ | 138.5 | 53 | 85.5 | $O_2$ | + |
| pJ2730 | $O_{sym}$ | 139.5 | 54 | 85.5 | $O_2$ | + |
| pJ2731 | $O_{sym}$ | 140.5 | 55 | 85.5 | $O_2$ | + |
| pJ2732 | $O_{sym}$ | 141.5 | 56 | 85.5 | $O_2$ | + |
| pJ2733 | $O_{sym}$ | 142.5 | 57 | 85.5 | $O_2$ | + |
| pJ2734 | $O_{sym}$ | 143.5 | 58 | 85.5 | $O_2$ | + |
| pJ2735 | $O_{sym}$ | 144.5 | 59 | 85.5 | $O_2$ | + |
| pJ2736 | $O_{sym}$ | 145.5 | 60 | 85.5 | $O_2$ | + |
| pJ2737 | $O_{sym}$ | 146.5 | 61 | 85.5 | $O_2$ | + |

*Denotes the base-pair distance (center to center) between the indicated elements

**Supplemental Table S3**. Series 2 spacing construct plasmids used in this study

| Plasmid | Distal Operator | Spacing (bp)* $O_{sym}\cdots O_2$ | $O_{sym}\cdots T_{sp}$ | $T_{sp}\cdots O_2$ | Proximal Operator | $T_{sp}$ |
|---|---|---|---|---|---|---|
| pJ2748 | $O_{sym}$ | 142.5 | 57 | 85.5 | $O_2$ | + |
| pJ2749 | $O_{sym}$ | 142.5 | 56 | 86.5 | $O_2$ | + |
| pJ2750 | $O_{sym}$ | 142.5 | 55 | 87.5 | $O_2$ | + |
| pJ2751 | $O_{sym}$ | 142.5 | 54 | 88.5 | $O_2$ | + |
| pJ2752 | $O_{sym}$ | 142.5 | 53 | 89.5 | $O_2$ | + |
| pJ2753 | $O_{sym}$ | 142.5 | 52 | 90.5 | $O_2$ | + |
| pJ2754 | $O_{sym}$ | 142.5 | 51 | 91.5 | $O_2$ | + |
| pJ2755 | $O_{sym}$ | 142.5 | 50 | 92.5 | $O_2$ | + |
| pJ2756 | $O_{sym}$ | 142.5 | 49 | 93.5 | $O_2$ | + |
| pJ2757 | $O_{sym}$ | 142.5 | 48 | 94.5 | $O_2$ | + |
| pJ2758 | $O_{sym}$ | 142.5 | 47 | 95.5 | $O_2$ | + |
| pJ2759 | $O_{sym}$ | 142.5 | 46 | 96.5 | $O_2$ | + |
| pJ2760 | $O_{sym}$ | 142.5 | 45 | 97.5 | $O_2$ | + |
| pJ2761 | $O_{sym}$ | 142.5 | 44 | 98.5 | $O_2$ | + |
| pJ2762 | $O_{sym}$ | 142.5 | 43 | 99.5 | $O_2$ | + |
| pJ2763 | $O_{sym}$ | 142.5 | 42 | 100.5 | $O_2$ | + |

*Denotes the base-pair distance (center to center) between the indicated elements

**Supplemental Table S4**. TALE and TALE-Nhp6A plasmids used in this study

| Plasmid | TALE | Nhp6A | LacI |
|---|---|---|---|
| pJ2657 | - | - | - |
| pJ2662 | - | - | + |
| pJ2658 | A (sp) | - | - |
| pJ2663 | A (sp) | - | + |
| pJ2660 | A (sp) | + | - |
| pJ2665 | A (sp) | + | + |
| pJ2741 | C (ns) | - | - |
| pJ2745 | C (ns) | - | + |
| pJ2743 | C (ns) | + | - |
| pJ2747 | C (ns) | + | + |

**Supplemental Table S7**. Protein-nucleic acid components of modeled TALE-DNA constructs.*

| Model | Amino-acid chain/residues | Base-pair chain/residues | |
|---|---|---|---|
| | A | B | C |
| 1 | 289-831 | 0-15 | 30-15 |
| 2 | 323-865 | 1-16 | 29-14 |
| 3 | 357-899 | 2-17 | 28-13 |
| 4 | 391-933 | 3-18 | 27-12 |
| 5 | 425-967 | 4-19 | 26-11 |
| 6 | 459-1001 | 5-20 | 25-10 |
| 7 | 493-1034 | 6-21 | 24-9 |
| 8 | 526-1068 | 7-22 | 23-8 |

*Chain identifiers and residue numbers of protein $C^{\alpha}$ and nucleic acid base atoms from pdb file 3ugm (1) incorporated in modeled 16-bp TALE-DNA constructs.

**Supplemental Table S8**. Rigid-body base-pair step parameters describing the average and specific closely related configurations of Nhp6A-bound DNA

| Base-pair step Average[*] | Rigid-body parameters ⟨Tilt⟩, deg | ⟨Roll⟩, deg | ⟨Twist⟩, deg | ⟨Shift⟩, Å | ⟨Slide⟩, Å | ⟨Rise⟩, Å |
|---|---|---|---|---|---|---|
| GG:CC | $-0.9_{\pm 1.8}$ | $-16.2_{\pm 5.7}$ | $29.4_{\pm 1.6}$ | $-0.91_{\pm 0.32}$ | $-0.75_{\pm 0.33}$ | $3.09_{\pm 0.08}$ |
| GT:AC | $4.3_{\pm 3.5}$ | $-9.3_{\pm 8.8}$ | $43.9_{\pm 2.8}$ | $1.43_{\pm 0.54}$ | $1.32_{\pm 0.37}$ | $3.85_{\pm 0.39}$ |
| TG:CA | $-4.3_{\pm 3.2}$ | $2.6_{\pm 6.0}$ | $22.6_{\pm 5.2}$ | $-0.03_{\pm 0.13}$ | $-0.23_{\pm 0.26}$ | $3.08_{\pm 0.37}$ |
| GA:TC | $0.7_{\pm 2.0}$ | $10.1_{\pm 1.6}$ | $25.4_{\pm 2.1}$ | $-0.05_{\pm 0.12}$ | $-1.77_{\pm 0.15}$ | $3.46_{\pm 0.19}$ |
| AT:AT | $-1.2_{\pm 1.9}$ | $-7.3_{\pm 5.1}$ | $28.8_{\pm 3.5}$ | $0.01_{\pm 0.18}$ | $-0.74_{\pm 0.17}$ | $3.78_{\pm 0.35}$ |
| TT:AA | $-2.8_{\pm 1.9}$ | $24.8_{\pm 6.1}$ | $25.2_{\pm 2.6}$ | $-0.45_{\pm 0.17}$ | $-0.31_{\pm 0.23}$ | $3.30_{\pm 0.22}$ |
| TG:CA | $0.5_{\pm 1.3}$ | $3.9_{\pm 4.9}$ | $23.0_{\pm 2.1}$ | $-1.07_{\pm 0.17}$ | $-1.36_{\pm 0.26}$ | $3.23_{\pm 0.13}$ |
| GT:AC | $2.5_{\pm 0.8}$ | $18.7_{\pm 2.9}$ | $34.0_{\pm 1.0}$ | $0.69_{\pm 0.19}$ | $0.23_{\pm 0.07}$ | $3.80_{\pm 0.10}$ |
| TT:AA | $-1.5_{\pm 0.8}$ | $19.0_{\pm 1.3}$ | $40.4_{\pm 0.3}$ | $0.08_{\pm 0.06}$ | $0.06_{\pm 0.04}$ | $3.06_{\pm 0.09}$ |

| Model 4[†] | ΔTilt, stdev | ΔRoll, stdev | ΔTwist, stdev | ΔShift, stdev | ΔSlide, stdev | ΔRise, stdev |
|---|---|---|---|---|---|---|
| GG:CC | 0.23 | 0.16 | 0.53 | 0.09 | 0.65 | 0.13 |
| GT:AC | 0.67 | 2.14 | 0.31 | 0.32 | 0.45 | 0.20 |
| TG:CA | 0.75 | 1.48 | 0.35 | 0.01 | 0.04 | 0.27 |
| GA:TC | 1.59 | 0.59 | 0.54 | 0.38 | 0.82 | 1.14 |
| AT:AT | 0.03 | 0.64 | 0.40 | 0.29 | 0.13 | 0.05 |
| TT:AA | 0.13 | 0.09 | 1.02 | 0.11 | 0.67 | 0.59 |
| TG:CA | 0.05 | 0.73 | 0.44 | 0.94 | 0.21 | 0.58 |
| GT:AC | 0.11 | 0.40 | 1.09 | 0.71 | 0.70 | 0.45 |
| TT:AA | 0.26 | 1.66 | 0.52 | 0.24 | 0.26 | 0.74 |

| Model 9[†] | ΔTilt, stdev | ΔRoll, stdev | ΔTwist, stdev | ΔShift, stdev | ΔSlide, stdev | ΔRise, stdev |
|---|---|---|---|---|---|---|
| GG:CC | 0.41 | 0.08 | 0.87 | 0.08 | 0.77 | 0.19 |
| GT:AC | 0.59 | 2.00 | 0.19 | 0.29 | 0.87 | 0.34 |
| TG:CA | 0.94 | 1.77 | 0.08 | 0.52 | 0.09 | 0.64 |
| GA:TC | 1.87 | 1.08 | 0.84 | 0.02 | 0.38 | 1.36 |
| AT:AT | 0.09 | 0.62 | 0.29 | 0.78 | 0.35 | 0.09 |
| TT:AA | 0.38 | 0.09 | 0.30 | 0.33 | 0.31 | 0.32 |
| TG:CA | 0.06 | 0.94 | 0.38 | 1.04 | 0.49 | 0.02 |
| GT:AC | 0.02 | 0.04 | 0.54 | 0.66 | 0.90 | 0.19 |
| TT:AA | 0.32 | 0.04 | 0.65 | 0.46 | 0.46 | 0.38 |

[*]Values at base-pair steps in direct contact with protein averaged over the 20 structures included in pdb file 1j5n (2).

[†]Deviation of base-pair step parameters in the specified model expressed relative to the average value and standard deviation of each parameter at the given step, i.e., $\Delta\theta_{XX:YY} = (\theta_{XX:YY} - \langle\theta_{XX:YY}\rangle)/\sigma_{XX:YY}$, where $\theta_{XX:YY}$ is one of the six parameters at base-pair step XX:YY, $\langle\theta_{XX:YY}\rangle$ is the average value of the parameter over all reported structures, and $\sigma_{XX:YY}$ is the corresponding standard deviation (stdev).

**Supplemental Table S9**. Properties of modeled loops bearing TALE and Nhp6A proteins.*

| $O_{sym}\cdots O_2$ spacing (bp) | TALE···Nh6pA spacing (bp) | Nh6pA orientation | Loop type | $J^{\dagger}$ | $C^{\alpha}\cdots C^{\alpha\ddagger}$ (Å) |
|---|---|---|---|---|---|
| 131.5 | 15 | f | $A_2^{F1}$ | $7.0\times10^{-5}$ | 23.8 |
| 131.5 | 21 | f | $A_1^{F1}$ | $8.9\times10^{-4}$ | 37.7 |
| 131.5 | 25 | r | $A_1^{F1}$ | $9.4\times10^{-4}$ | 53.1 |
| 137.5 | 15 | f | $P_1^{F1}$ | $6.5\times10^{-8}$ | 22.9 |
| 137.5 | 21 | f | $P_2^{F2}$ | $9.7\times10^{-7}$ | 38.9 |
| 137.5 | 25 | r | $P_2^{F2}$ | $6.3\times10^{-7}$ | 56.8 |

*Loops from Fig. 6 and Supplemental Fig. S7 with proportions greater than 10%
†Estimated $J$ factors of loops of the specified types.
‡Distances between the $C^{\alpha}$ atoms at the N terminus of the TALE protein and the C terminus of Nhp6A in the dominant configurations of the loops illustrated in Fig. 6 and Supplemental Fig. S7. Protein-free DNA modeled with a 10.7-bp helical repeat. Note the altered distances between chain ends upon re-orientation of Nhp6A.

### Supplemental references

1. Mak, A.N., Bradley, P., Cernadas, R.A., Bogdanove, A.J. and Stoddard, B.L. (2012) The crystal structure of TAL effector PthXo1 bound to its DNA target. *Science*, **335**, 716-719.
2. Masse, J.E., Wong, B., Yen, Y.M., Allain, F.H., Johnson, R.C. and Feigon, J. (2002) The S. cerevisiae architectural HMGB protein NHP6A complexed with DNA: DNA and protein conformational changes upon binding. *J Mol Biol*, **323**, 263-284.
3. Swigon, D., Coleman, B.D. and Olson, W.K. (2006) Modeling the Lac repressor-operator assembly: the influence of DNA looping on Lac repressor conformation. *Proc Natl Acad Sci U S A*, **103**, 9879-9884.
4. Swigon, D. and Olson, W.K. (2008) Mesoscale modeling of multi-protein-DNA assemblies: the role of the catabolic activator protein in Lac-repressor-mediated looping. *Int J Non Linear Mech*, **43**, 1082-1093.
5. Bell, C.E. and Lewis, M. (2000) A closer view of the conformation of the Lac repressor bound to operator. *Nat Struct Biol*, **7**, 209-214.
6. Lewis, M., Chang, G., Horton, N.C., Kercher, M.A., Pace, H.C., Schumacher, M.A., Brennan, R.G. and Lu, P. (1996) Crystal structure of the lactose operon repressor and its complexes with DNA and inducer. *Science*, **271**, 1247-1254.
7. Spronk, C.A., Bonvin, A.M., Radha, P.K., Melacini, G., Boelens, R. and Kaptein, R. (1999) The solution structure of Lac repressor headpiece 62 complexed to a symmetrical lac operator. *Structure*, **7**, 1483-1492.
8. Romanuka, J., Folkers, G.E., Biris, N., Tishchenko, E., Wienk, H., Bonvin, A.M., Kaptein, R. and Boelens, R. (2009) Specificity and affinity of Lac repressor for the auxiliary operators O2 and O3 are explained by the structures of their protein-DNA complexes. *J Mol Biol*, **390**, 478-489.
9. Britton, L.A., Olson, W.K. and Tobias, I. (2009) Two perspectives on the twist of DNA. *J Chem Phys*, **131**, 245101.
10. Clauvelin, N., Olson, W.K. and Tobias, I. (2012) Characterization of the geometry and topology of DNA pictured as a discrete collection of atoms. *J Chem Theory Comput*, **8**, 1092-1107.
11. Lu, X.J. and Olson, W.K. (2008) 3DNA: a versatile, integrated software system for the analysis, rebuilding and visualization of three-dimensional nucleic-acid structures. *Nat Protoc*, **3**, 1213-1227.
12. Lu, X.J., Bussemaker, H.J. and Olson, W.K. (2015) DSSR: an integrated software tool for dissecting the spatial structure of RNA. *Nucleic Acids Res*, **43**, e142.
